# Low-Dose Interleukin-2 Resolves Stress-Induced Chronic Sensitization Independent of Opioid Receptor Signaling

**DOI:** 10.64898/2026.09.01.748528

**Authors:** Róli R. Simões, Scheila Kraus, Jintao Zhang, Leandro Flores do Nascimento, Dilip K. Tosh, Jacqueline Unsinger, Richard S. Hotchkiss, Tao Che, Kenneth A. Jacobson, Yu-Qing Cao

## Abstract

Stress is a common trigger of headache and widespread somatic pain. Repeated exposure to stress establishes latent sensitization that is tonically suppressed by endogenous inhibitory systems. Using brain-penetrating and peripherally restricted receptor antagonists in a mouse model of repetitive restraint stress, we systematically identified these protective pathways. Peripheral kappa opioid receptors as well as GABA type A and type B (GABA_A_ and GABA_B_) receptors, alongside central opioid and cannabinoid signaling, actively suppress headache-related facial mechanical hypersensitivity in stressed mice. Blocking any pathway quickly reinstated cephalic allodynia. Peripheral opioid and GABA_A_ receptor signaling also inhibited stress-induced latent sensitization on hindpaw. In contrast, depletion of anti-inflammatory regulatory T (Treg) cells prolonged facial but not hindpaw sensitization, suggesting that Tregs preferentially limit stress-induced chronic headache. Next, we administered low-dose interleukin-2 (LD-IL-2) to stressed mice to preferentially expand and activate Treg cells. Following LD-IL-2 treatment, neither subthreshold pain triggers nor blockade of endogenous inhibitory pathways reinstated cephalic or hindpaw allodynia in either sex, indicating elimination of stress-induced sensitization. Mechanistically, LD-IL-2 acted through Treg cells to recruit multiple peripheral cytokine pathways without engaging endogenous opioid, GABA, or cannabinoid receptor signaling. Notably, LD-IL-2 was more effective than anti-CGRP (calcitonin gene-related peptide) therapy in preventing headache-related chronic sensitization in stressed mice. Collectively, these findings reveal multiple central and peripheral pathways that act concertedly to mask stress-induced latent sensitization and strongly support further evaluation of LD-IL-2 as a novel treatment for stress-induced headache and widespread pain with a distinct mechanism of action.

## 1. Introduction

Stress and its relief are common triggers for migraine and widespread somatic pain, which frequently co-occur and cause profound disability [94]. Repeated exposure to stress promotes a state of latent sensitization and increases the risk of pain episodes during the interictal (pain-free) period [28; 78]. Identifying the endogenous mechanisms that maintain the interictal stability is critical for developing more effective treatments.

In rodent models, repetitive restraint stress induces acute facial and hindpaw mechanical hypersensitivity that transitions into the latent sensitization state [4; 42; 67; 77]. Primary afferent neurons exhibit hyperalgesic priming and exaggerated responses to otherwise subthreshold stimuli [4; 42; 69]. The hyperexcitable pain circuits are actively suppressed by the constitutive activation of endogenous inhibitory pathways [4; 42; 67; 77], including opioid peptides acting through mu, delta, and kappa opioid receptors (MOR, DOR, and KOR); gamma-aminobutyric acid (GABA) acting through type A and B (GABA_A_ and GABA_B_) receptors; and endocannabinoids acting through type-1 and type-2 (CB_1_ and CB_2_) receptors. Disrupting these pathways can rapidly unmask latent sensitization, revealing the precarious equilibrium that maintains the pain-free state [5; 15; 19; 37; 56; 60; 63; 81; 84-86; 90]. Although many studies have focused on central mechanisms [5; 15; 37; 60; 63; 81; 84; 86; 90], peripheral inhibitory pathways also suppress latent sensitization [11; 22; 27; 32; 50; 56; 89]. After repetitive stress, endogenous enkephalin reduces migraine-like behavior through peripheral DOR, but not MOR [56], whereas amygdalar KOR signaling promotes stress-induced pain behaviors [42; 87; 88]. Both central and peripheral cannabinoid pathways attenuate sensitization in other headache models [85; 89]. However, it is not clear whether they ameliorate stress-induced headache, as chronic stress can disrupt endocannabinoid tone [30; 34]. Whether peripheral KOR or GABAergic signaling suppresses stress-induced latent sensitization remains largely unexplored.

Beyond neurochemical pathways, immune cells also contribute to pain resolution [31]. In addition to maintaining immune tolerance and constraining inflammation, endogenous regulatory T (Treg) cells limit behavioral sensitization in models of post-traumatic headache and neuropathic pain [16; 25; 45; 58; 59; 92]. Repetitive stress, however, has produced inconsistent effects on circulating and splenic Treg cells in rodent models [9; 35; 40], and their role in stress-induced headache and body pain is unknown. Treg-enhancing strategies attenuate chronic pain-related behaviors across multiple mouse models through context-dependent mechanisms, including secretion of anti-inflammatory cytokines such as transforming growth factor-β1 (TGF-β1), interleukin-10 (IL-10), interleukin-35, and release of enkephalins that activate DOR [3; 16; 18; 23; 25; 33; 36; 48; 58; 59; 91; 92]. Low-dose interleukin-2 (LD-IL-2) treatment, which preferentially expands and activates Treg cells [41; 76], prevents chronic headache-related sensitization in non-stress models [16; 91]. Whether it prevents stress-induced chronic sensitization and which pathways mediate its effects remain unclear. Here, using a mouse model of repetitive restraint stress that recapitulates key clinical features of migraine and somatic pain [4; 42], we systematically identified peripheral inhibitory systems that mask latent sensitization, evaluated the efficacy of LD-IL-2 against a current migraine-preventive strategy, and delineated its mechanisms of action (MoA) across Treg dependence, cytokine signaling, and canonical neurochemical pathways.

## 2. Materials and methods

### 2.1. Mice

All experiments reported in this study were conducted in strict accordance with the recommendations in the Guide for the Care and Use of Laboratory Animals of the National Institutes of Health and were approved by the Institutional Animal Care and Use Committee at Washington University in St. Louis (approval no. 25-0178). To avoid social isolation stress, all mice were group housed (2-5 per cage, same sex) in the animal facility of Washington University in St. Louis on a 12-hour light-dark cycle with constant temperature (23-24°C), humidity (45-50%), and *ad libitum* access to standard laboratory diet and water. All experiments were performed during the light phase of cycle (9 am – 5 pm). Female and male CD-1 mice (8 – 12 weeks old, 25 – 35 g) were purchased from Charles River (O’Fallon, MO).

Adult C57BL/6J mice and DEREG (depletion of regulatory T cell, 32050) breeders on C57BL/6J background were purchased from the Jackson Laboratory (Bar Harbor, ME). The DEREG mice contain a transgenic allele that expresses the diphtheria toxin receptor-enhanced green fluorescent protein (DTR-EGFP) fusion protein under the control of the genomic sequences that regulate the expression of endogenous *Foxp3* [43]. DEREG breeders were crossed with C57BL/6J mice to generate heterozygotes for experiments. The genotype was determined by polymerase chain reaction (PCR) of tail DNA as described [43].

### 2.2. Repetitive restraint stress paradigm

Mice were subjected to 2 hours of restraint stress in the morning for 3 consecutive days as detailed in previous studies [4; 42; 72]. Briefly, mice were guided into individual cylindrical rodent restrainers (Stoelting 51338) facing the acrylic front. The tail was threaded through the moveable disk, and the disk was tightened so the mouse was incapable of movement but still maintained normal respiration. Mice were checked every 5 minutes (min) to ensure that they did not change the position or suffer from any injury. Sham mice were kept in their home cages in a different room, without access to water or food during the same period.

### 2.3. Bright light stress (BLS) paradigm

Mice were placed in a clear acrylic chamber (11 x 11 x 15 cm) surrounded with LED light strips. The center light intensity was 1000 lux. Freely moving mice were exposed to bright light for 15 min to induce stress [62]. Facial mechanical responses were assessed 1 hour after exposure to the bright light. The BLS paradigm did not elicit behavioral responses in sham mice.

### 2.4. Behavioral tests

Mice were randomly distributed between the experimental groups, extensively handled by the experimenters, and were well-habituated to the test room and the test apparatus. The experimenters were blinded to the treatment groups during data collection and analysis.

#### 2.4.1. Withdrawal thresholds to mechanical stimuli on facial skin

One to three days before testing, mice were lightly anesthetized with isoflurane. The hair on the mouse forehead (above and between the eyes) was shaved and trimmed with a Multi-Groom Ultra Precise Beard Styler trimmer Groomer (Philips, PHMG1100/16, Cambridge MA). On the test day, the experimenters gently held the mouse on the palm with minimal restraint and applied a series of calibrated von Frey filaments perpendicularly to the shaved skin, causing the filaments to bend for 5 seconds. A positive response was determined by the following criteria as in previous studies: eye squinting, stroking face with the forepaw, head withdrawal from the stimulus, or head shaking [73; 91]. Dixon’s up-down paradigm was used to determine the 50% withdrawal threshold [10].

After the facial mechanical threshold returned to baseline level, mice were challenged with various stimuli to assess chronic sensitization. Facial mechanical thresholds were measured before and 10 min to 3 hours after the stimuli.

#### 2.4.2. Facial pain score

To measure the aversiveness to facial mechanical stimuli, the responses of mice to individual von Frey filaments (0.07 g, 0.16 g and 0.40 g) were scored based on the most aversive behavior they display [16; 72]: 0) no responses; 1) detection: eye squinting, facial skin crinkling and/or a small head shaking; 2) withdrawal: head withdrawal from the filament and/or forepaw stroking of face once; 3) vigorous forepaw stroking of face 2 or more times; and 4) vigorous head shaking and/or attacking the filament. Scores 1 and 2 represented mostly reflexive behaviors, and scores 3 and 4 represented affective-motivational behaviors that reflect the unpleasantness of pain [8; 14; 16; 72]. For each mouse at each time point, a cumulative pain score was generated by combining the 9 individual scores.

#### 2.4.3. Facial grimace score

Facial grimace score was quantified as described in previous studies to assess ongoing pain [4; 16; 44]. Mice were habituated in individual clear acrylic boxes (10 x 10 x 10 cm). After being exposed to a subthreshold stimulus, mice were recorded by a video camera for 40 min. Individual frames were obtained every 2 min during the last 30 min of the video. Within each 2-min recording, the first clear, unobstructed head shot was grabbed for analysis. Behaviors on five portions of the face (orbital tightening, nose bulge, check bulge, flattening of ears, and flattening of whiskers) were scored on a scale of 0 – 2 (0 = not present, 1 = high confidence of a moderate appearance or equivocation of present or not, 2 = high confidence of a severe appearance present). For each mouse, a grimace score was generated by averaging scores from all portions of the face from all snapshots. Each frame was analyzed by two experimenters, and the scores were averaged to assess ongoing pain.

#### 2.4.4. Withdrawal thresholds to punctate mechanical stimuli on hindpaw skin

Mice were acclimated for 30 min in individual clear acrylic cylinder (11 x 11 x 15 cm) on an elevated wire mesh (6 x 6 mm) platform to allow access to the glabrous skin of the hindpaw. A series of calibrated von Frey filaments (0.07 – 2.0 g) were used to apply mechanical stimuli to the plantar surface of the hindpaw as previously described [36]. A response was defined as the mouse lifting and/or shaking its paw from the filament. A median paw withdrawal threshold was determined using an adaption of the Dixon’s up-and-down method [10]. The thresholds of both paws were averaged to yield a single value for individual datapoints.

Responses to mechanical stimuli on the hindpaw were measured before and after repetitive restraint stress. After the mechanical thresholds returned to the baseline level, mice received various stimuli to evaluate chronic sensitization. Hindpaw mechanical thresholds were measured before and 3 hours after the stimuli. Note that after intraplantar injection of prostaglandin E2 (PGE_2_), the withdrawal thresholds were only measured in the injected paw.

#### 2.4.5. Responses to dynamic mechanical stimuli on hindpaw skin

Mice were acclimated for 20 min in individual clear acrylic cylinder on an elevated wire mesh platform. The medial plantar surface of the hind paw was stimulated by light stroking with a blunt paintbrush (5/0, Princeton Art & Brush Co., Princeton, NJ, USA) from heel to toe at ∼2 cm/second. The behavior was scored as described previously [36]: 0) no evoked movement or lifting the stimulated paw for less than 1 second; 1) a sustained lifting (more than 2 seconds) of the stimulated paw toward the body or a single gentle flinching of the stimulated paw; 2) a strong lateral lift of the stimulated paw above the level of the body or a startle-like jump; 3) multiple flinching or licking of the stimulated paw. Each paw was tested 3 times, with minimal interval of 3 min. The allodynia scores of both paws were averaged to indicate the response to dynamic mechanical stimuli for individual mice. However, after intraplantar injection of PGE_2_, the allodynia scores were only measured in the injected paw.

### 2.5. Drug administration

#### 2.5.1. Sodium nitroprusside (SNP)

The nitric oxide donor SNP (Sigma, St. Louis, MO) was freshly diluted from the stock (200 mg/ml in saline at 4°C) with saline before intraperitoneal (i.p.) injection to mice in a dose of 0.3 mg/kg [72]. In one experiment testing the effects of LD-IL-2, mice received i.p. injection of 1 mg/kg SNP. Behavioral responses were assessed 3 hours after SNP administration.

#### 2.5.2. PGE_2_

PGE_2_ was freshly diluted from the stock solution (Sigma, 1 µg/µL in saline, aliquoted and stored at -20°C). After the recovery of stress-induced hind paw hypersensitivity, mice received intraplantar injection of PGE_2_ (100 ng in 20 µl saline) and were tested 3 hours later to assess hyperalgesic priming [38; 65]. In one experiment testing the effects of LD-IL-2, mice received intraplatar injection of 300 ng PGE_2_.

#### 2.5.3. Opioid receptor antagonists

The pan-opioid receptor antagonist naloxone (Sigma) was freshly diluted from the stock solution (50 mg/ml in saline at 4°C) before i.p. injection to mice in a dose of 2 mg/kg. The peripherally-acting, MOR and KOR-preferred antagonist methylnaltrexone bromide (MNB, Sigma) was freshly diluted from the stock (1 mg/ml in water at -20°C) with saline before i.p. injection to mice in a dose of 2 mg/kg [6; 26]. The DOR antagonist naltrindole (Sigma) was freshly diluted from the stock solution (8 mg/ml in water at -20°C) before subcutaneous injection to mice in a dose of 3 mg/kg. A peripherally-acting, DOR-preferred antagonist TAN-452 (MedChemExpress, Monmouth Junction, NJ) was freshly diluted from the stock solution (10 mg/ml in 10% dimethyl sulfoxide [DMSO], 40% PEG 300, 5% Tween 80 and 45% saline at -20°C) before s.c. injection to mice in a dose of 10 mg/kg [80]. The peripherally-acting KOR antagonist MRS7299 was freshly diluted from the stock solution (5 mg/ml in DMSO at 4°C) before i.p. injection to mice at 0.003 – 0.1 mg/kg [82]. Behavioral responses were assessed 1 – 3 hours after the drug administration.

#### 2.5.4. GABA receptor antagonists

The peripherally-acting GABA_A_ receptor antagonist bicuculline methiodide (Bicuc, Sigma) was freshly diluted from the stock solution (10 mg/ml in saline at 4°C) before i.p. injection to mice in a dose of 3 mg/kg [52; 75]. The peripherally-acting GABA_B_ receptor antagonist 2-OH-saclofen (Sac, Sigma) was freshly diluted from the stock solution (0.5 mg/ml in water at -20°C) before i.p. injection to mice in a dose of 2 mg/kg [39; 51; 83]. Behavioral responses were assessed 10 min (for Sac) and 3 hours (for Bicuc) after the drug administration.

#### 2.5.5. Cannabinoid receptor antagonists

The peripherally restricted CB_1_ receptor antagonist AM6545 as well as the brain-penetrating CB_1_ receptor antagonist AM281 and CB_2_ receptor antagonist AM630 (all from Cayman Chemical, Ann Arbor, MI) were freshly diluted from the stock solution (3 – 10 mg/ml in DMSO at -20°C) before i.p. injection to mice in a dose of 10 mg/kg for AM6545 [1], 0.5 mg/kg for AM281 [7; 12] and 1 mg/kg for AM630 [49], respectively. Behavioral responses were assessed 3 hours after the drug administration.

#### 2.5.6. Olcegepant/BIBN4096BS (BIBN)

Twelve days after the repetitive stress, when acute sensitization was fully resolved, the small molecule calcitonin gene-related peptide (CGRP) receptor antagonist BIBN (Sigma, 4 nmol in 20 µl saline) was injected subcutaneously under the periorbital skin 0.5 hour before and 2.5 hour after SNP administration, respectively [53]. Facial mechanical thresholds were measured 0.5 and 2.5 hours after BIBN injection.

#### 2.5.7. Depletion of Treg cells

Diphtheria toxin (DT, Sigma) was freshly diluted from the stock (2 mg/ml in saline, aliquoted and stored at -80°C) before each administration. To deplete the endogenous Treg cells, DEREG mice received i.p. injections of 0.5 µg DT in 100 µl saline for two consecutive days every 5 days [16; 54].

#### 2.5.8. LD-IL-2 treatment

Recombinant mouse IL-2 (carrier-free, Biolegend, San Diego, CA) was freshly diluted from the stock (0.87 µg/µL aliquoted and stored at -80°C) daily. Mice received daily i.p. injections of saline or LD-IL-2 (1 µg in 100 µL of saline) for 5 – 9 times. Note that LD-IL-2 was always administered after the stress induction or the completion of behavioral tests.

#### 2.5.9. Neutralizing antibodies and control immunoglobulin G (IgG)

Neutralizing antibodies and control IgGs were purchased from BioXcell (Lebanon, NH), stored at 4°C, and i.p. administered (200 µg/mouse) after stress induction or the completion of behavioral tests on the same day. The anti-interferon-γ (IFN-γ) antibody (BE0055) and control IgG (BE0088) were injected every 3 days.[13] The anti-IL-10 antibody (BE0049) and control IgG (BE0088) were injected every 2 days [33; 46]. The neutralizing antibody against all TGF-β isoforms (BE0057) and control IgG (BE0083) was injected every 3 days [33; 47].

The chimeric recombinant anti-CGRP antibody was generated by adapting the V_H_ and V_L_ sequences (from IMGT/mAb-DB) of the CGRP neutralizing antibody Fremanezumab to mouse IgG1(κ) backbone. The antibody was produced by Genscript in HD CHO-S cells and purified by MabSelect SuRe LX affinity chromatography with > 80% purity and < 1.3 EU/mg endotoxin as in our previous study.[72] The anti-CGRP antibody and control IgG (BE0083) were stored at 4°C and i.p. administered at 30 mg/kg after the resolution of stress-induced acute sensitization.

### 2.6. Blood collection, antibody staining and flow cytometry

Mice used for flow cytometry analysis were not used in behavioral tests. DEREG mice were subjected to sham or repetitive restraint stress procedure for 3 consecutive days. About 100 µl blood was collected from each mouse by submandibular bleeding at each timepoint. After the last blood collection, mice were euthanized and spleens were collected, ground and filtered through a sterile 70 µm cell strainer. After lysis of red blood cells (420301; Biolegend), cells were pelleted, resuspended, and stained with the antibodies that recognize mouse CD3ε (clone 145-2C11), CD4 (clone GK1.5), and CD25 (clone PC61, all from Biolegend) for 20 min in the darkness. Flow cytometry data were collected with FACScan (Becton Dickinson, Franklin Lakes, NJ). CellQuest Pro (Becton Dickinson) and Rainbow X Alias (Cytek, Fremont, CA) were used for data acquisition and fluorescence compensation. The frequencies of individual cell subpopulations were determined with the FlowJo software (v10.8.1, Becton Dickinson).

### 2.7. Tissue preparation, immunohistochemistry, and image analysis

Immunohistochemistry and image analysis were performed as previously described [91], with experimenters blinded to the treatment groups during data collection and analysis. Briefly, DEREG mice were euthanized with i.p. injection of barbiturate (200 mg/kg) and were transcardially perfused with warm 0.1 M phosphate buffered saline (PBS, pH 7.2) followed by cold 4% formaldehyde in 0.1 M phosphate buffer (PB, pH 7.2) for fixation. Lumbar L4 dorsal root ganglion (DRG) tissues were collected and post-fixed for 4 hours. After incubating the DRG in 0.1 M PB with 30% sucrose for 2-3 days, tissues were embedded in OCT (optimal cutting temperature) compound and sectioned at 15 µm on a cryostat and collected on Superfrost Plus glass slides in sequence.

One in every 3 DRG sections were processed for immunohistochemistry. The sections were dried at room temperature, washed three times in 0.01 M PBS, and incubated in blocking buffer consisting 0.01 M PBS, 10% normal goat serum (NGS), and 0.3% triton X-100 for 1 hour at RT. Sections were then incubated overnight with the chicken anti-EGFP antibody (1:1000 dilution in blocking buffer, AVES Lab, Davis, CA) in a humidity chamber at 4°C overnight. After 6 washes (5 min each) in washing buffer containing 0.01 M PBS with 1% NGS and 0.3% triton and 3 rinses in 0.01 M PBS, sections were incubated with blocking buffer for 1 hour, followed by the incubation with AlexaFluor 488-conjugated secondary antibody (1:1000 dilution in blocking buffer, Invitrogen, Waltham, MA) at room temperature for 1 hour. After washing off the antibodies, sections were rinsed with PBS, cover-slipped using Fluoromount-G Slide Mounting Medium (Electron Microscopy, Hatfield, PA), sealed with nail polish, and stored at 4°C.

Immunofluorescence was observed through a 40x objective on a Nikon TE2000S inverted epifluorescence microscope. To quantify Treg cells in DRG, all EGFP^+^ cells in individual sections were counted, and the number was multiplied by 3 to obtain the total number of cells per ganglion in each mouse.

### 2.8. Statistical analysis

For behavioral experiments, power analysis was conducted to estimate sample size with > 80% power to reach a significance level of 0.05. All data were reported as mean ± standard error of the mean (SEM). The Shapiro-Wilk test was used to check data normality. Statistical significance between experimental groups was assessed using Graph Pad software (Graph Pad Prism, 9.5 version, San Diego, CA). Two-tailed t-test, one-way or two-way repeated-measures analysis of variance (RM-ANOVA) with post hoc Bonferroni test for multiple comparison were used where appropriate. Differences with *P* < 0.05 were considered statistically significant. The statistical analysis for individual experiments was described in figure legends and Supplementary Table 1.

## 3. Results

### 3.1. Repetitive restraint stress engages multiple endogenous inhibitory pathways to suppress headache-related chronic sensitization

First, we characterized repetitive restraint stress-induced behaviors that are mechanistically related to migraine headache in our experimental setting. Female CD-1 mice were subjected to 2-hour restraint sessions over three consecutive days as in previous studies [4; 42]. This resulted in a week-long reduction in the withdrawal thresholds to mechanical stimuli on periorbital skin, indicating the development of headache-related facial skin hypersensitivity (Fig. 1A). After the resolution of acute sensitization, we challenged mice with migraine triggers SNP (0.3 mg/kg, i.p.) and BLS (1000 lux, 15 min). Sham mice did not respond to SNP or BLS, confirming that they are subthreshold stimuli under basal condition. In stressed mice, both SNP and BLS quickly re-established facial mechanical hypersensitivity (Fig. 1B), revealing the development of hyperalgesic priming (hyper-responsiveness to subthreshold stimuli) as shown in previous studies [4; 42].

**Figure 1.**
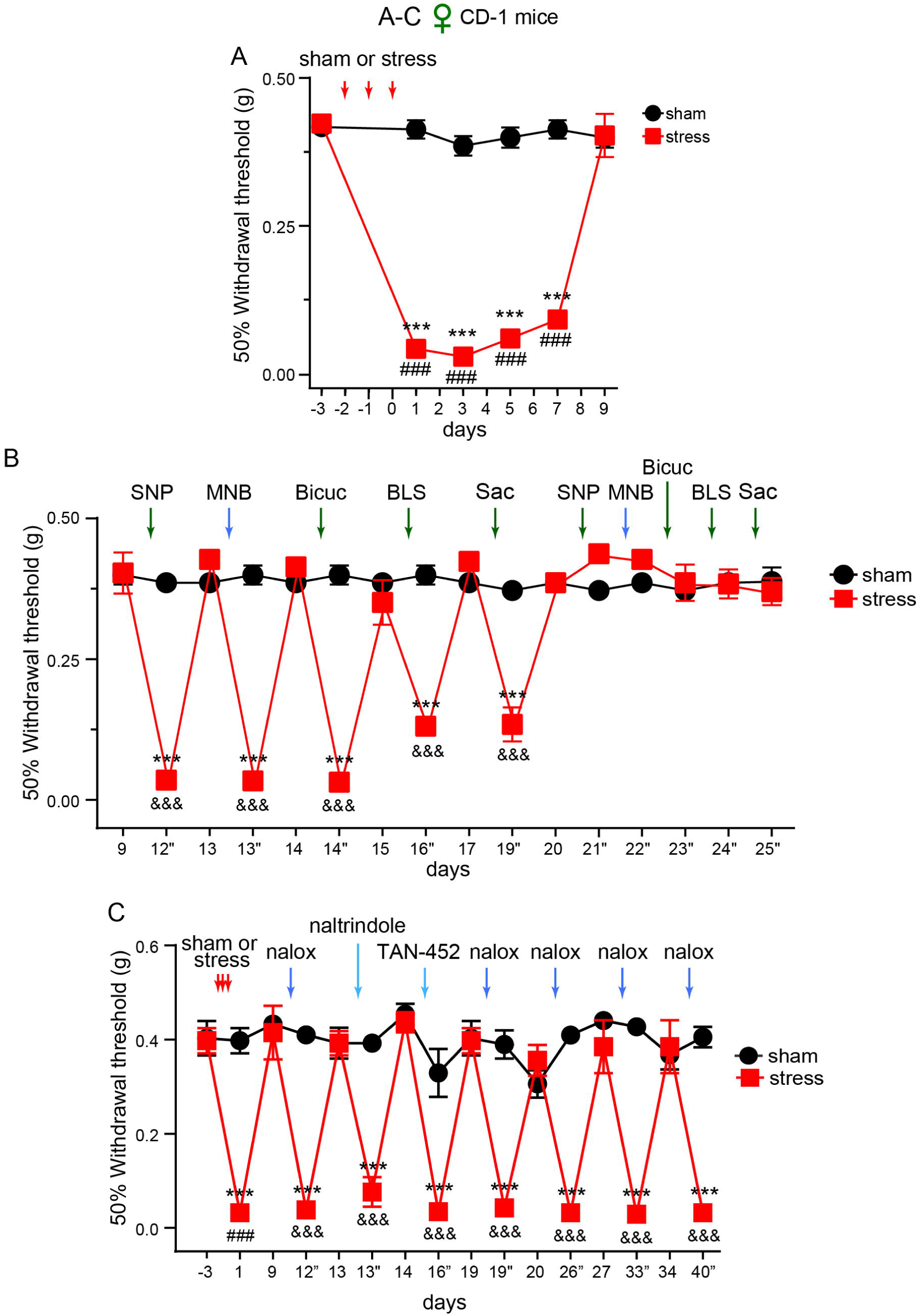
Repetitive restraint stress engages multiple endogenous inhibitory pathways to suppress headache-related chronic sensitization. **(A)** Repetitive stress decreased the facial withdrawal threshold in female CD-1 mice (*n* = 6/group). \*\*\**P* < 0.001, sham versus stress groups; ^###^*P* < 0.001, compared with the baseline thresholds on day -3 in the stress group. **(B)** Repetitive stress resulted in a masked state of heightened sensitivity after the resolution of acute sensitization (same mice as in **A**) as revealed by the i.p. administration of sodium nitroprusside (SNP, 0.3 mg/kg), methylnaltrexone bromide (MNB, 2 mg/kg), bicuculline methiodide (Bicuc, 3 mg/kg), 2-OH-saclofen (Sac, 2 mg/kg) or exposure to bright light stress (BLS). By day 21-25, neither stimulus could evoke facial skin hypersensitivity in stressed mice. Timepoints followed by ” indicate that the thresholds were measured 10 minutes to 3 hours after the administration of drugs or exposure to BLS. \*\*\**P* < 0.001, sham versus stress groups; ^&&&^*P* < 0.001, relative to the thresholds before SNP, MNB, Bicuc, BLS, or Sac challenges in the stress group. **(C)** Naloxone (nalox, 2 mg/kg, i.p.), naltrindole (3 mg/kg, s.c.) and TAN-452 (10 mg/kg, i.p.) administration unmasked facial mechanical hypersensitivity 2-6 weeks post-stress in female CD-1 mice (n = 6/group). \*\*\**P* < 0.001, sham versus stress groups; ^###^*P* < 0.001, compared with the day -3 baseline thresholds in the stress group. ^&&&^*P* < 0.001, relative to the thresholds before each naloxone or naltrindole administration in the stress group.

To investigate whether repetitive stress induces latent sensitization, mice were treated with various peripherally-acting drugs to block the endogenous inhibitory pathways, including the opioid receptor antagonist MNB (2 mg/kg, i.p.), GABA_A_ receptor selective antagonist Bicuc (3 mg/kg, i.p.), and GABA_B_ receptor antagonist Sac (2 mg/kg, i.p.). Sham mice did not respond to any of the drugs. On the contrary, all three antagonists elicited robust facial skin hypersensitivity in stressed mice (Fig. 1B), indicating that peripheral opioid, GABA_A_ and GABA_B_ signaling pathways actively suppress latent sensitization. The chronic sensitization phase lasted for about 2 weeks (Fig. 1B). By day 21-25 post-stress, none of the subthreshold stimuli could re-establish facial mechanical hypersensitivity (Fig. 1B). These results indicate that repetitive restraint stress engages multiple endogenous inhibitory pathways to mask headache-related chronic sensitization.

To gain more insight into how endogenous opioid receptor signaling regulates stress-induced sensitization, we treated mice with naloxone (2 mg/kg, i.p.) to block all opioid receptor subtypes in peripheral and brain tissues. Notably, naloxone still evoked facial skin hypersensitivity 4-6 weeks after repetitive stress (Fig. 1C, day 26, 33 and 40), when the peripherally-acting MNB was no longer effective (Fig. 1B), indicating that central opioid receptors continue to mask stress-induced sensitization after the disengagement of peripheral opioid signaling. We also investigated the contribution of individual opioid receptor subtypes. Consistent with previous studies [21; 56; 66], subcutaneous administration of DOR-antagonist naltrindole (3 mg/kg) reinstated facial skin hypersensitivity (Fig. 1C), indicating that the signaling through DOR inhibits latent sensitization. Treating mice with a peripherally-acting, DOR-preferring antagonist TAN-452 (10 mg/kg, i.p. [80]) also resulted in facial skin hypersensitivity (Fig. 1C). However, whether the dose of TAN-452 we used selectively block peripheral DOR signaling remains to be determined.

MNB preferentially inhibits peripheral MOR and KOR [6; 26]. Since MOR antagonist does not evoke behavioral sensitization in this stress paradigm [56], we investigated whether MNB-induced facial skin hypersensitivity results from inhibition of peripheral KOR. To test this hypothesis, mice received i.p. injection of MRS7299, a peripherally-acting, KOR-selective full antagonist [82]. At 0.1 mg/kg, MRS7299 significantly reduced the facial mechanical thresholds in stressed mice but not in sham controls, whereas vehicle injection had no effect (Fig. 2A). We conclude that peripheral KOR is activated by repetitive stress to suppress chronic sensitization.

**Figure 2.**
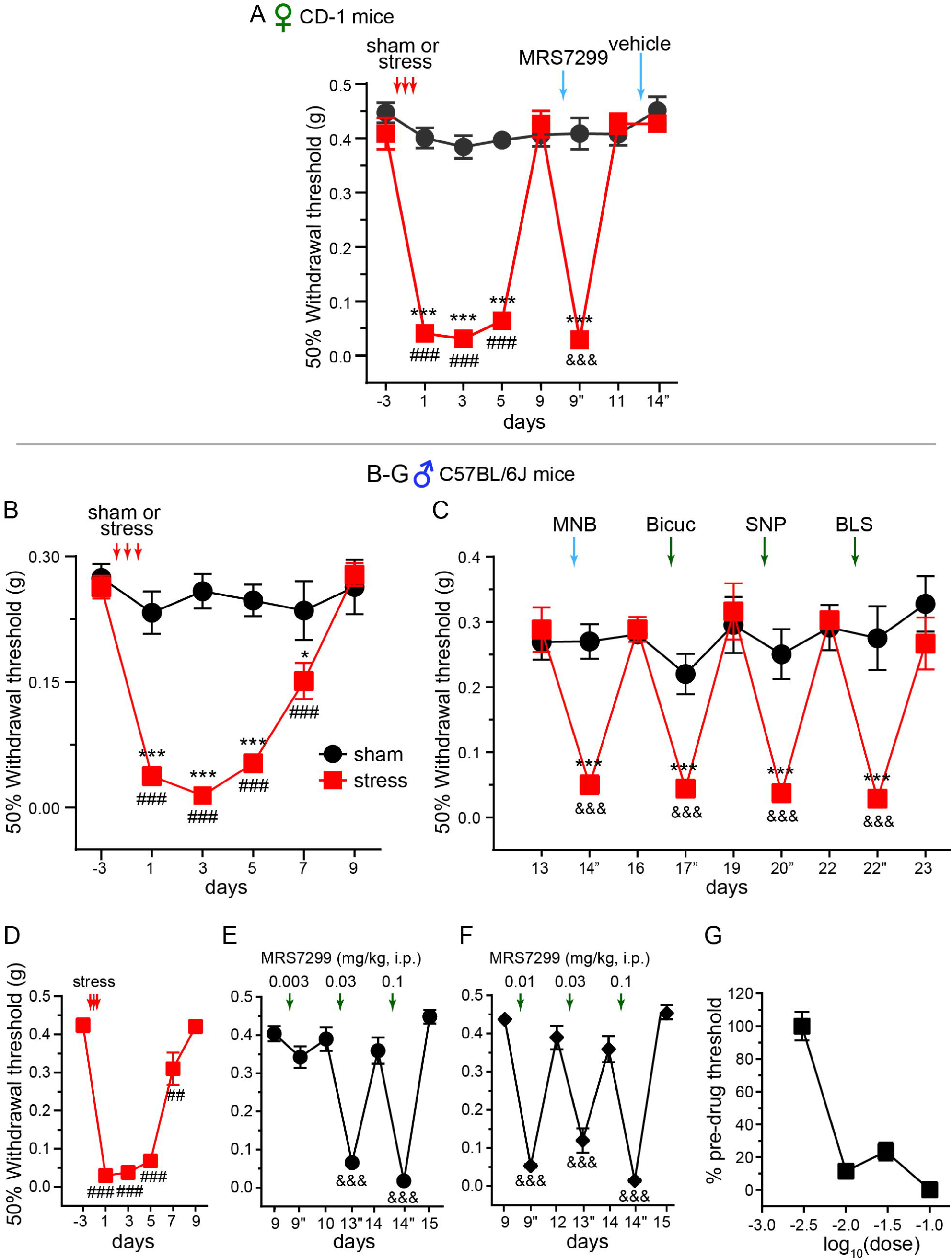
Repetitive restraint stress activates peripheral KOR in both male and female mice. **(A)** MRS7299 administration (0.1 mg/kg, i.p.) evoked robust facial skin hypersensitivity in stressed female CD-1 mice but not sham controls (n = 8/group). Injection of the vehicle for MRS7299 did not alter facial mechanical responses in either group. \*\*\**P* < 0.001, sham versus stress groups; ^###^*P* < 0.001, compared with the day -3 baseline thresholds in the stress group. ^&&&^*P* < 0.001, relative to the thresholds before MRS7299 administration in the stress group. (**B**) Repetitive stress resulted in similar magnitude and duration of acute sensitization in male C57BL/6J mice (*n* = 5-6/group). \**P* < 0.05, \*\*\**P* < 0.001, sham versus stress groups; ^###^*P* < 0.001, compared with the baseline thresholds on day -3 in the stress group. (**C**) Repetitive stress-induced chronic sensitization in male C57BL/6J mice (same as in **E**) was revealed by the administration of MNB, Bicuc, SNP and exposure to BLS. \*\*\**P* < 0.001, sham versus stress groups; ^&&&^*P* < 0.001, relative to the thresholds before SNP, MNB, Bicuc, BLS, or Sac challenges in the stress group. **(D-G)** Dose-response study of MRS7299 in male C57BL/6J mice. **(C)** Repetitive stress-induced acute sensitization was confirmed in all mice (*n* = 10), ^##^*P* < 0.01, ^###^*P* < 0.001, compared with the baseline thresholds on day -3. (**E-F)** Mice were then divided into two groups (*n* = 5/group) to receive various doses of MRS7299, ^&&&^*P* < 0.001, relative to the thresholds before MRS7299 administration. **(G)** Dose-response curve of MRS7299 in male C57BL/6J mice. Facial mechanical thresholds after MRS7299 administration were normalized to the corresponding pre-drug values.

To address the potential sex-and strain-dependence of stress-induced sensitization, we applied the same repetitive restraint stress paradigm to male C57BL/6J mice and observed comparable acute sensitization as in female outbred CD-1 mice (Fig. 1A and 2B). After the resolution of acute phase, administration of MNB, Bicuc, SNP or exposure to BLS all quickly re-established facial skin hypersensitivity (Fig. 2C). Latent sensitization could also be induced by 0.01-0.1 mg/kg of MRS7299, confirming the activation of peripheral KOR in stressed male C57BL/6J mice (Fig. 2D-G). Taken together, these findings indicate that repetitive stress engages multiple endogenous inhibitory pathways to suppress chronic sensitization in both male and female, inbred and outbred mice.

### 3.2. Peripheral opioid and GABA_A_ receptor signaling pathways inhibit repetitive stress-induced chronic somatic sensitization

Stress or its relief also triggers widespread body pain. Indeed, the same repetitive restraint stress paradigm also caused a significant reduction of the withdrawal threshold to punctate mechanical stimuli (von Frey filaments) on hindpaw skin in both male and female CD-1 mice (Fig. 3A and Supplementary Fig. 1, day 1-4). After the mechanical thresholds returned to the baseline level, intraplantar injection of 100 ng PGE_2_ re-established the hindpaw hypersensitivity to punctate stimuli (Fig. 3A, day 6”; Supplementary Fig. 1, day 7” and 8”), indicating the development of hyperalgesic priming. By day 9 post-stress, PGE_2_ could no longer induce hindpaw hypersensitivity (Supplementary Fig. 1). Thus, relative to the headache-related facial skin hypersensitivity, repetitive restraint stress resulted in a much shorter duration of somatic sensitization. Sham mice did not show changes of hindpaw mechanical responses throughout the experiment (Fig. 3A, sham group).

**Figure 3.**
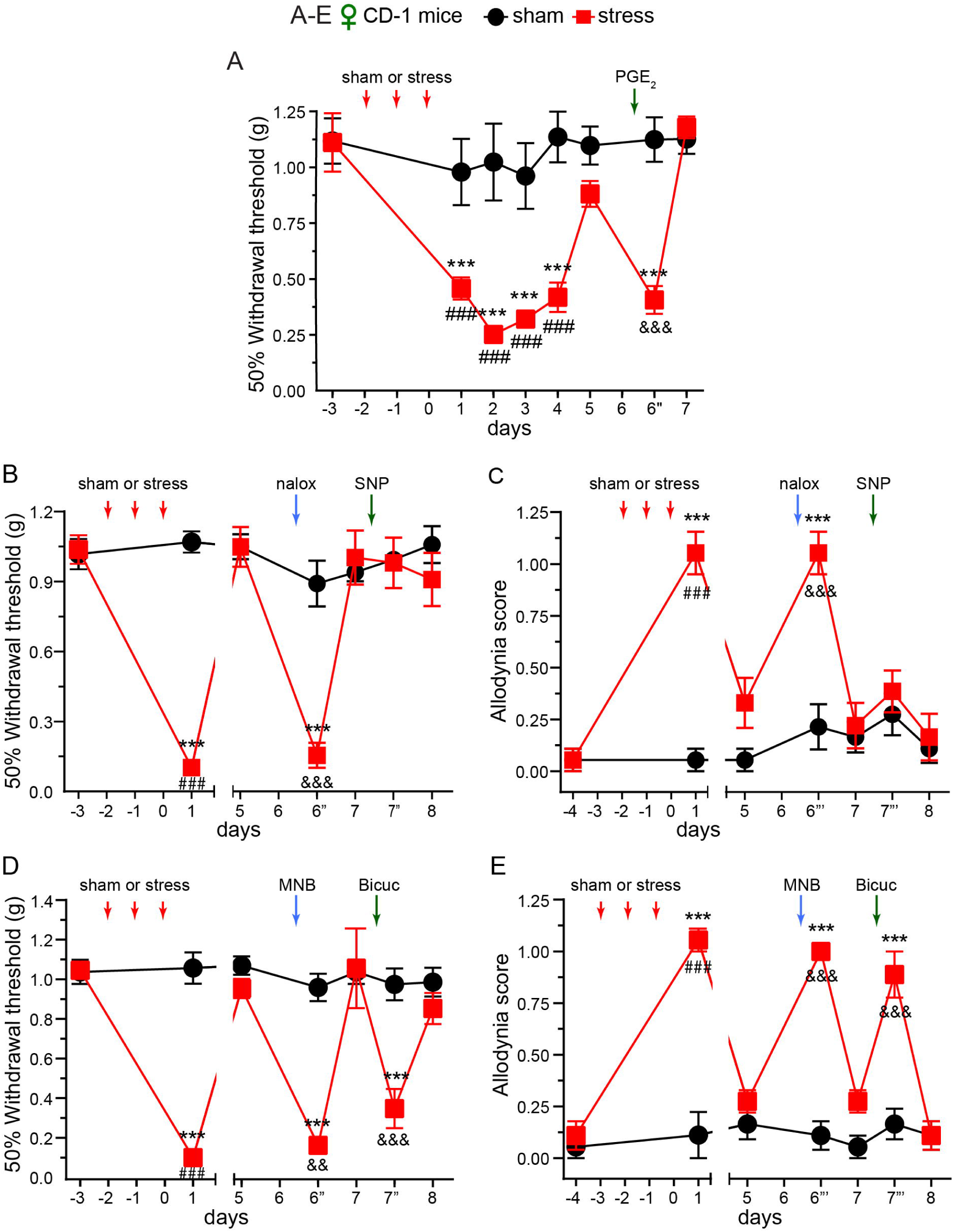
Peripheral opioid and GABA_A_ receptor signaling pathways inhibit repetitive stress-induced chronic somatic sensitization. **(A)** Repetitive restraint stress resulted in behavioral sensitization related to widespread pain. acute sensitization and hyperalgesic priming of the hindpaw glabrous skin in female CD-1 mice (*n* = 4-8/group). \*\**P* < 0.01, \*\*\**P* < 0.001, sham versus stress groups. Day 1-4, stress-induced acute sensitization, ^###^*P* < 0.001, compared with the day -3 baseline thresholds in the stress group. Day 6”, stress-induced hyperalgesic priming revealed by intraplantar injection of 100 ng Prostaglandin E2 (PGE_2_). ^&&&^*P* < 0.001, relative to the day 5 thresholds in the stress group. **(B-C)** Systemic injection of naloxone revealed repetitive stress-induced latent sensitization to punctate **(B)** and dynamic (**C)** mechanical stimuli on the hindpaw of female CD-1 mice (*n* = 6/group). Timepoints followed by ”’ indicate that the dynamic allodynia was scored after measuring the withdrawal thresholds in the same mice. \*\*\**P* < 0.001, sham versus stress groups; ^###^*P* < 0.001, compared with the pre-stress baseline values in the stress group; ^&&&^*P* < 0.001, relative to the day 5 values baseline thresholds in the stress group. **(D-E)** Both systemic MNB and Bicuc administration disinhibited repetitive stress-induced latent sensitization to punctate **(D)** and dynamic **(E)** mechanical stimuli on the hindpaw of female CD-1 mice (*n* = 6/group). \*\*\**P* < 0.001, sham versus stress groups; ^###^*P* < 0.001, compared with the pre-stress baseline values in the stress group; ^&&&^*P* < 0.001, relative to the values before MNB and Bicuc administration in the stress group.

In subsequent experiments (Fig. 3B-E), we found that repetitive restraint stress resulted in hypersensitivity to both punctate (Fig. 3B and 3D) and dynamic (brush, Fig. 3C and 3E) mechanical stimuli on the hindpaw skin of female CD-1 mice. After the resolution of acute sensitization, hindpaw skin hypersensitivity could be reinstated by the administration of naloxone, MNB or Bicuc in stressed mice but not in sham control (Fig. 3B-E), indicating that activation of both peripheral opioid and GABA_A_ receptor signaling pathways masks repetitive stress-induced somatic latent sensitization.

The presence of stress-induced hindpaw skin sensitization raises the concern that the facial skin hypersensitivity in stressed mice may result from sensitization of the facial skin afferents rather than the headache-related meningeal afferents. To test this possibility, we selectively blocked the sensitization of facial skin afferents as in a previous study [53] by injecting olcegepant (BIBN4096BS, 4 nmol in 20 µl saline), the small molecule CGRP receptor antagonist under the periorbital skin of stressed mice (Fig. 4A). Although systemic administration of a neutralizing antibody against CGRP prevents SNP-induced facial skin hypersensitivity in stressed mice [4], periorbital injection of olcegepant did not prevent SNP from reinstating facial skin sensitization (Fig. 4B). Moreover, systemic injection of SNP re-established mechanical hypersensitivity on facial but not hindpaw skin (Fig. 1B, 2C versus Fig. 3B-C). This is in line with the fact that infusion of nitric oxide donor triggers migraine-like headache but not other somatic pain in migraine patients [79]. Together, these results support the notion that repetitive stress-induced facial skin hypersensitivity is mechanistically related to headache rather than resulting from the sensitization of facial skin afferents.

**Figure 4.**
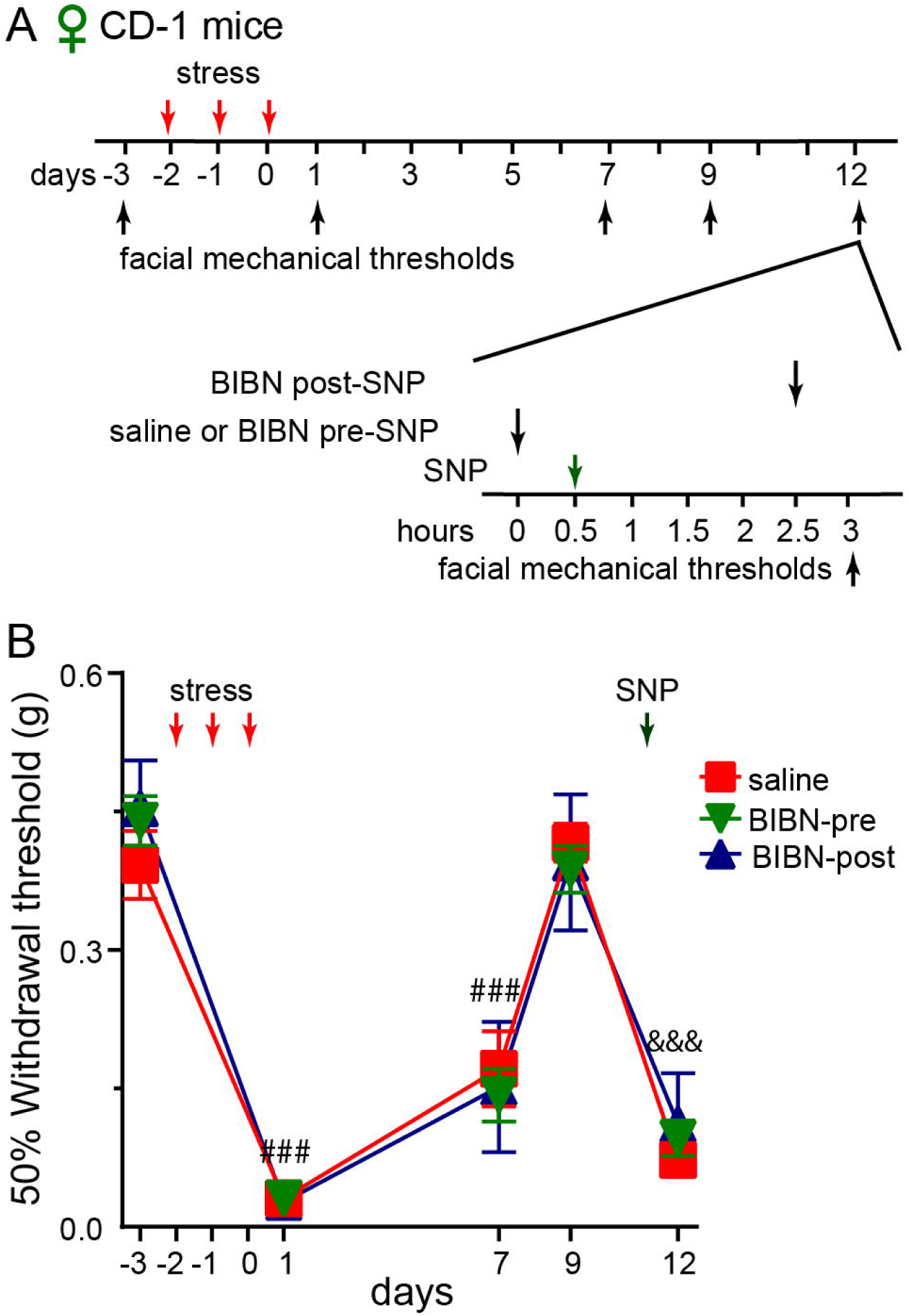
Blocking CGRP signaling in periorbital skin does not prevent SNP-induced re-sensitization in stressed mice. **(A)** Timeline of the experiment. BIBN: CGRP receptor antagonist BIBN4096BS. **(B)** Periorbital injection of BIBN (4 nmol in 20 µl saline) did not inhibit repetitive stress-induced hyperalgesic priming in female CD-1 mice (*n* = 4-6/group). ^###^*P* < 0.001, compared with the day -3 baseline threshold within individual groups. ^&&&^*P* < 0.001, compared with the day 9 thresholds within individual groups.

### 3.3. The anti-inflammatory Treg cells differentially regulate headache-and somatic pain-related sensitization in stressed mice

In addition to opioid and GABA_A_ receptor signaling, the endogenous Treg cells facilitate the resolution of post-traumatic headache-related behaviors in our previous work [16]. Using DEREG mice that selectively express DTR-EGFP in Treg cells [43], we first examined whether repetitive stress changes the abundance of Tregs and other T cells by flow cytometry (Fig. 5A). We used EGFP and CD25 as markers to identify Treg cells and saw comparable levels of EGFP^+^CD4^+^ and CD25^+^CD4^+^ cells in the peripheral blood of sham and stressed mice (Fig. 5B-C). The abundance of total circulating T cells (CD3^+^) and the CD4^+^ subset was not altered by stress either (Fig. 5D-E). The frequencies of EGFP^+^CD4^+^ and CD25^+^CD4^+^ Tregs as well as CD3^+^ and CD4^+^ T cells in the spleen were also comparable between sham and stress groups (Fig. 5F-I), indicating that the number of Treg cells is not affected by this repetitive stress paradigm.

**Figure 5.**
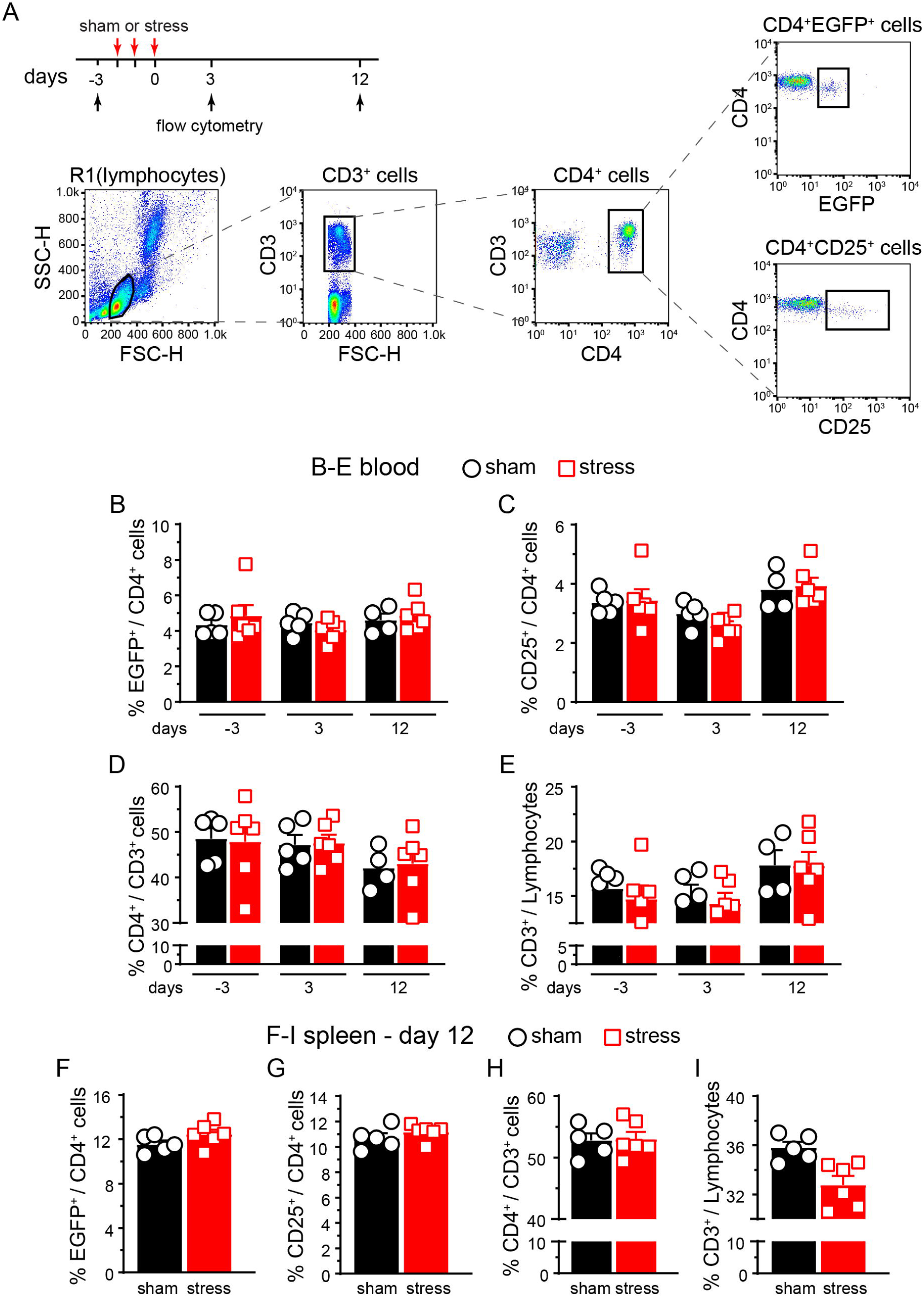
Repetitive restraint stress does not alter the abundance of Treg cells. **(A)** Timeline of the experiment and gating strategy for individual T cell subpopulations. **(B-E)** The frequencies of EGFP^+^ **(B)** and CD25^+^ **(C)** cells among CD4^+^ T cells, CD4^+^ cells among CD3^+^ T cells **(D)** and CD3^+^ T cells **(E)** in the peripheral blood of sham and stressed male DEREG mice (*n* = 5-6/group). **(F-I)** The frequencies of EGFP^+^CD4^+^ **(F)**, CD25^+^CD4^+^ **(G)**, CD4^+^ **(H)** and CD3^+^ **(I)** T cells in the spleen of sham and stressed male DEREG mice (same as in **B-E**).

Next, we investigated whether endogenous Treg cells regulates stress-induced sensitization. Male and female DEREG mice on C57BL/6J background received 0.5 µg DT for two consecutive days every 5 days to deplete the endogenous Treg cells for about 2 weeks [16; 54]. Stress-induced acute sensitization and hyperalgesic priming on facial skin were comparable between wild-type and saline-treated DEREG mice (Fig. 1-2 and 6A). In DT-treated DEREG mice, the resolution of acute sensitization was significantly delayed (Fig. 6A, day 9-14). During the priming phase, SNP-induced facial skin hypersensitivity resolved within 24 hours in mice with intact Treg cells (Fig. 6A, saline group, day 14”-15) but persisted for at least 7 days in DT-treated DEREG mice (Fig. 6A, DT group, day 14”-21). We also scored the responses to weak, mild, and strong mechanical stimuli on facial skin to monitor the aversiveness/unpleasantness of pain [8; 14; 16; 72]. One day after SNP administration, the facial pain score was significantly higher in DT-treated DEREG mice relative to the saline control (Fig. 6B, day 15). By day 27 post-stress, SNP did not evoke facial skin hypersensitivity in either saline-or DT-treated DEREG mice (Fig. 6A). These results indicate that endogenous Treg cells limit the persistence of headache-related sensitization and reduce the aversiveness of pain in response to repetitive stress.

**Figure 6.**
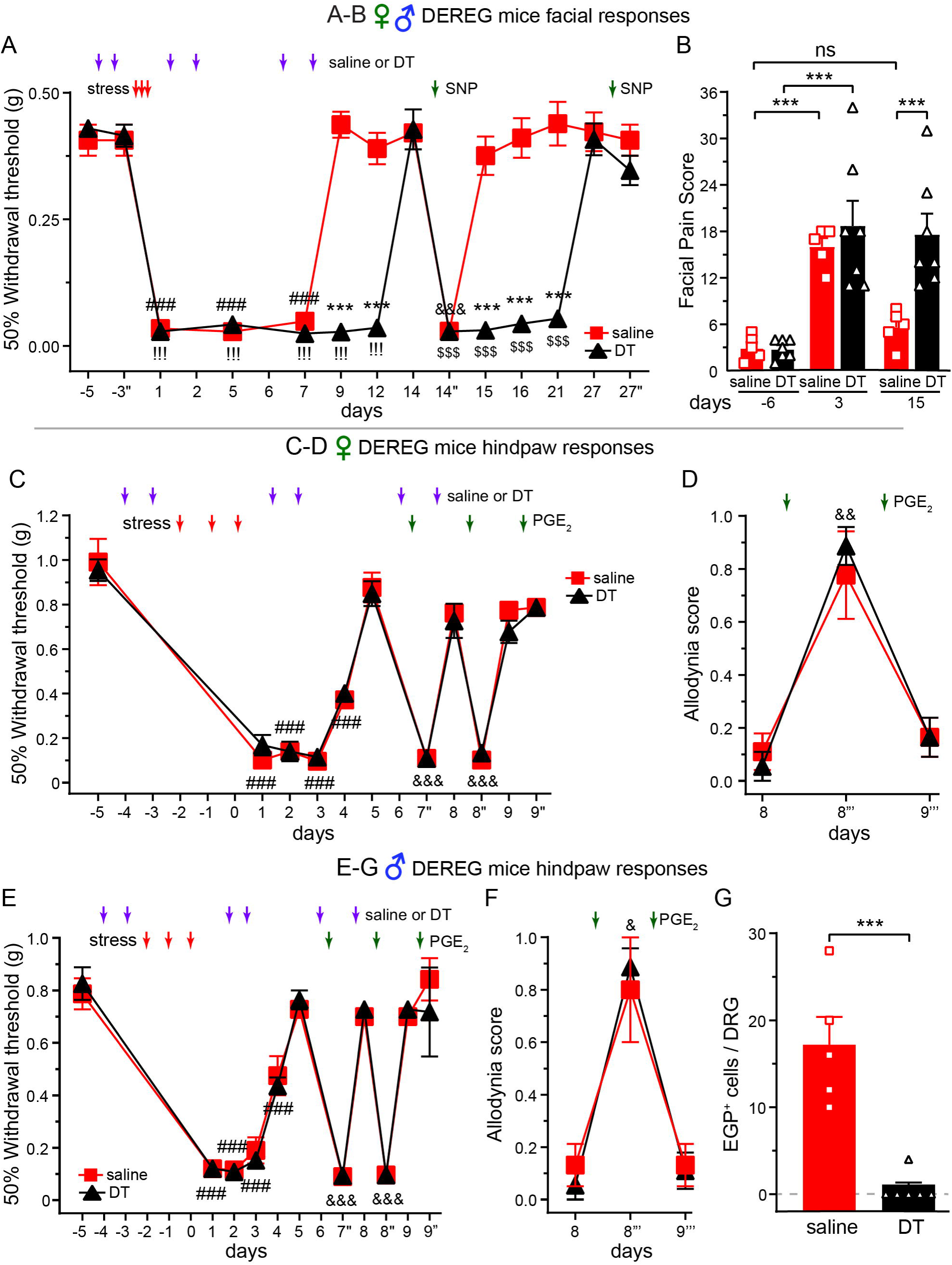
Endogenous Treg cells selectively inhibit headache-related sensitization after repetitive stress. **(A)** Repeated diphtheria toxin (DT, 0.5 µg/mouse/day, i.p.) administration prolonged stress-induced acute and chronic sensitization in both male and female DEREG mice (*n* = 3-4 male and 2-3 female mice/group). \*\*\**P* < 0.001, saline versus DT groups; ^###^*P* < 0.001, ^!!!^*P* < 0.001, compared with the pre-stress baseline thresholds in saline and DT groups, respectively; ^&&&^*P* < 0.001, ^$$$^*P* < 0.001, relative to the day 14 thresholds in saline and DT groups, respectively. **(B)** Treg depletion prolonged SNP-induced increase in facial pain score (same mice as in **A**). \*\*\**P* < 0.001. **(C, E)** Repeated DT administration did not alter stress-induced hindpaw sensitization to punctate mechanical stimuli in female (**C**, *n* = 6/group) and male DEREG mice (**E**, *n* = 5-6/group). ^###^*P* < 0.001, compared with the pre-stress baseline thresholds within individual groups; ^&&&^*P* < 0.001, relative to the day 14 thresholds within individual groups. **(D, F)** Treg depletion did not affect stress-induced hindpaw dynamic allodynia in DEREG mice (same as in **C** and **E**, respectively). ^#^*P* < 0.05, ^##^*P* < 0.01, compared with the day 8 allodynia scores within individual groups. (**G**) Repeated DT administration effectively depleted EGFP^+^ Treg cells in L4 DRG of DEREG mice (same as in **E**). Samples were collected 1 day after the last behavioral assay. \*\*\**P* < 0.001.

We went on to examine whether Treg cells suppress stress-induced cutaneous allodynia on hindpaw. Surprisingly, repeated DT administration did not alter stress-induced sensitization to either punctate or dynamic mechanical stimuli on hindpaw skin in male or female DEREG mice (Fig. 6C-F, DT versus saline groups). In both saline-and DT-treated mice, stress-induced acute sensitization resolved by day 5 and hyperalgesic priming persisted till day 8, as indicated by the responses to repeated hindpaw injections of PGE_2_ (Fig. 6C, E, day 5-9”). After the behavioral assay, we quantified the number of EGFP^+^ Treg cells in the lumbar L4 DRG of DEREG males on day 9. DRG from DT-treated mice exhibited a substantial reduction (> 90%) of Treg cells relative to the saline control (Fig. 6G), indicating that repeated DT administration effectively depleted Treg cells in peripheral tissues. Collectively, these data identify a previously unrecognized dissociation between stress-induced headache and body pain, with endogenous Treg cells selectively inhibit facial but not hindpaw sensitization after repetitive stress.

### 3.4. LD-IL-2 treatment alleviates repetitive stress-induced hindpaw skin sensitization

To test the effectiveness of Treg-targeted therapy on stress-induced body pain, we gave female CD-1 mice 1-week of daily LD-IL-2 treatment (1 µg/mouse/day, i.p.) during and after repetitive stress (Fig. 7A, day -2 to day 4). Saline-treated mice developed robust hindpaw hypersensitivity and hyperalgesic priming. LD-IL-2 treatment prevented stress-induced acute sensitization (Fig. 7A, day 1-4). Three days after the cessation of LD-IL-2, when the mechanical threshold returned to baseline level, intraplantar injection of PGE_2_ only slightly reduced the mechanical threshold (Fig. 7A, day 7’’), indicating that stress-induced hyperalgesic priming is substantially attenuated. These data support the therapeutic effect of early LD-IL-2 treatment on stress-induced widespread pain.

**Figure 7.**
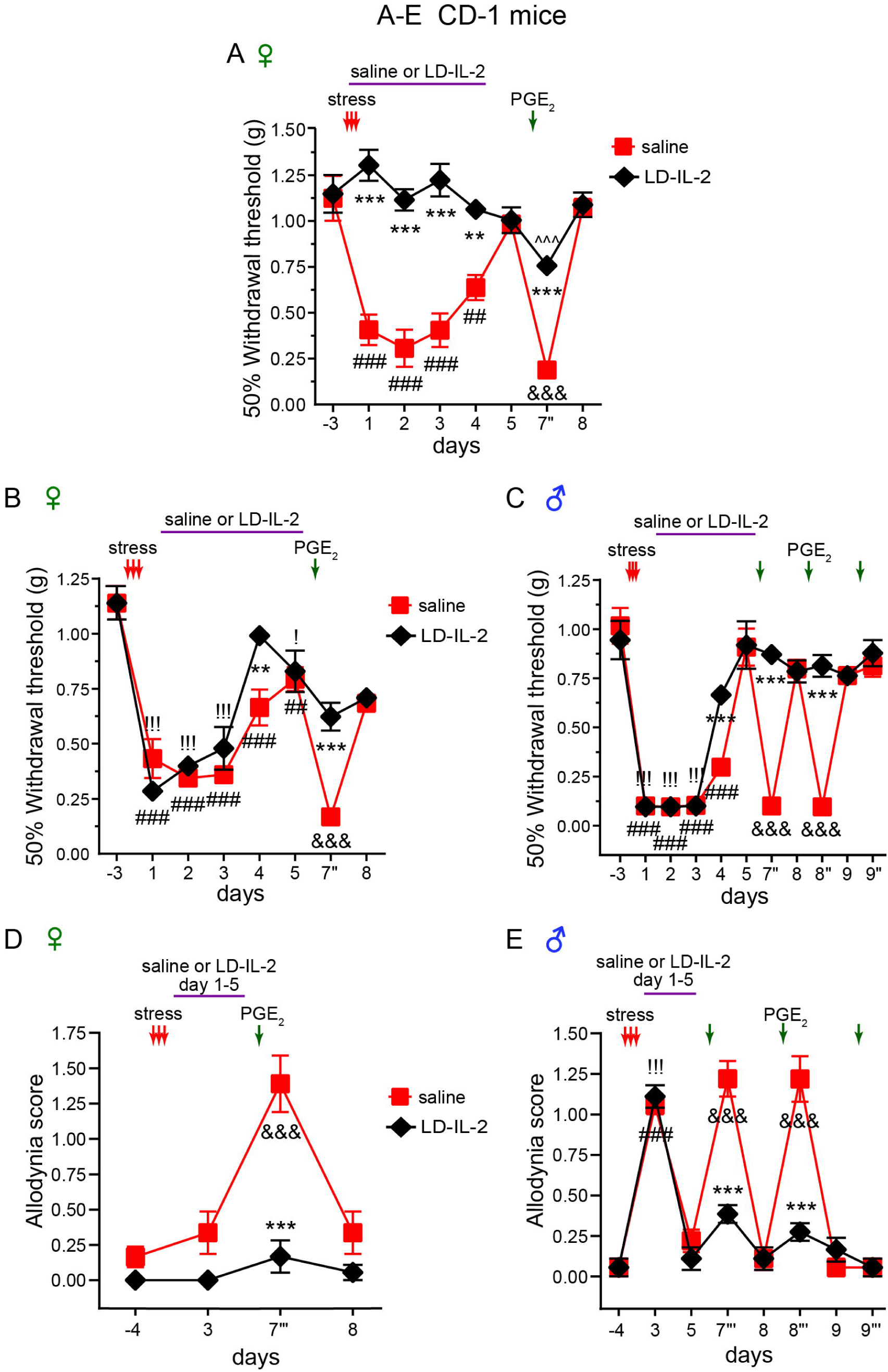
LD-IL-2 treatment prevents PGE_2_-induced hindpaw mechanical allodynia in stressed mice. **(A)** Pretreatment with LD-IL-2 (1 µg/mouse/day, i.p.) prevented stress-induced hindpaw mechanical hypersensitivity and substantially reduced hyperalgesic priming in female CD-1 mice (*n* = 5-6/group). \*\**P* < 0.01, \*\*\**P* < 0.001, saline versus LD-IL-2 groups; ^##^*P* < 0.01, ^###^*P* < 0.001, compared with the baseline threshold on day -3; ^&&&^*P* < 0.001, ^^^^^*P* < 0.01, compared with the day 5 thresholds before PGE_2_ injection in saline and LD-IL-2 groups, respectively. **(B-C)** LD-IL-2 facilitated the resolution of repetitive stress-induced acute hindpaw skin sensitization and prevented hyperalgesic priming to punctate mechanical stimuli in female **(B)** and male **(C)** CD-1 mice (*n* = 6/group). \*\**P* < 0.01, \*\*\**P* < 0.001, saline versus LD-IL-2 groups; ^##^*P* < 0.01, ^###^*P* < 0.001, compared with the day -3 baseline threshold in the saline group; ^!^*P* < 0.05, ^!!!^*P* < 0.001, compared with the day -3 baseline threshold in the LD-IL-2 group; ^&&&^*P* < 0.001, compared with the values before each PGE_2_ injection in the saline group. **(D-E)** Post-stress LD-IL-2 prevented PGE_2_-induced dynamic allodynia in stressed female **(D)** and male **(E)** CD-1 mice (same as in Fig. 4B**-C**). \*\*\**P* < 0.001, saline versus LD-IL-2 groups; ^###^*P* < 0.001, ^!!!^*P* < 0.001, compared with the day -4 baseline scores in saline and LD-IL-2 groups, respectively; ^&&&^*P* < 0.001, relative to the values before each PGE_2_ injection in the saline group.

To be more translationally relevant, we started LD-IL-2 treatment in both male and female CD-1 mice 1 day after repetitive stress, when the acute sensitization was already established (Fig. 7B-C, day 1-5). The magnitude and duration of stress-induced sensitization were comparable in male and female mice (Fig. 7B-C, saline groups). LD-IL-2 treatment accelerated the resolution of acute sensitization (Fig. 7B-C, day 4) and prevented PGE_2_-induced hindpaw punctate allodynia in both sexes (Fig. 7B-C, day 5-9). The hypersensitivity to dynamic mechanical stimuli after intraplantar PGE_2_ was also abolished by LD-IL-2 treatment (Fig. 7D-E).

In subsequent experiments, we assessed the effects of LD-IL-2 on stress-induced activation of endogenous opioid and GABA_A_ receptors. After the resolution of acute sensitization, both male and female mice exhibited naloxone-and Bicuc-evoked hindpaw hypersensitivity to punctate and dynamic mechanical stimuli (Fig. 8A-D, saline groups). This latent sensitization was absent in LD-IL-2 treated mice (Fig. 8A-D, LD-IL-2 groups). These results suggest that post-stress LD-IL-2 treatment can prevent stress-induced hyperalgesic priming and latent sensitization, thereby blocking the transition from acute to chronic widespread pain in both males and females.

**Figure 8.**
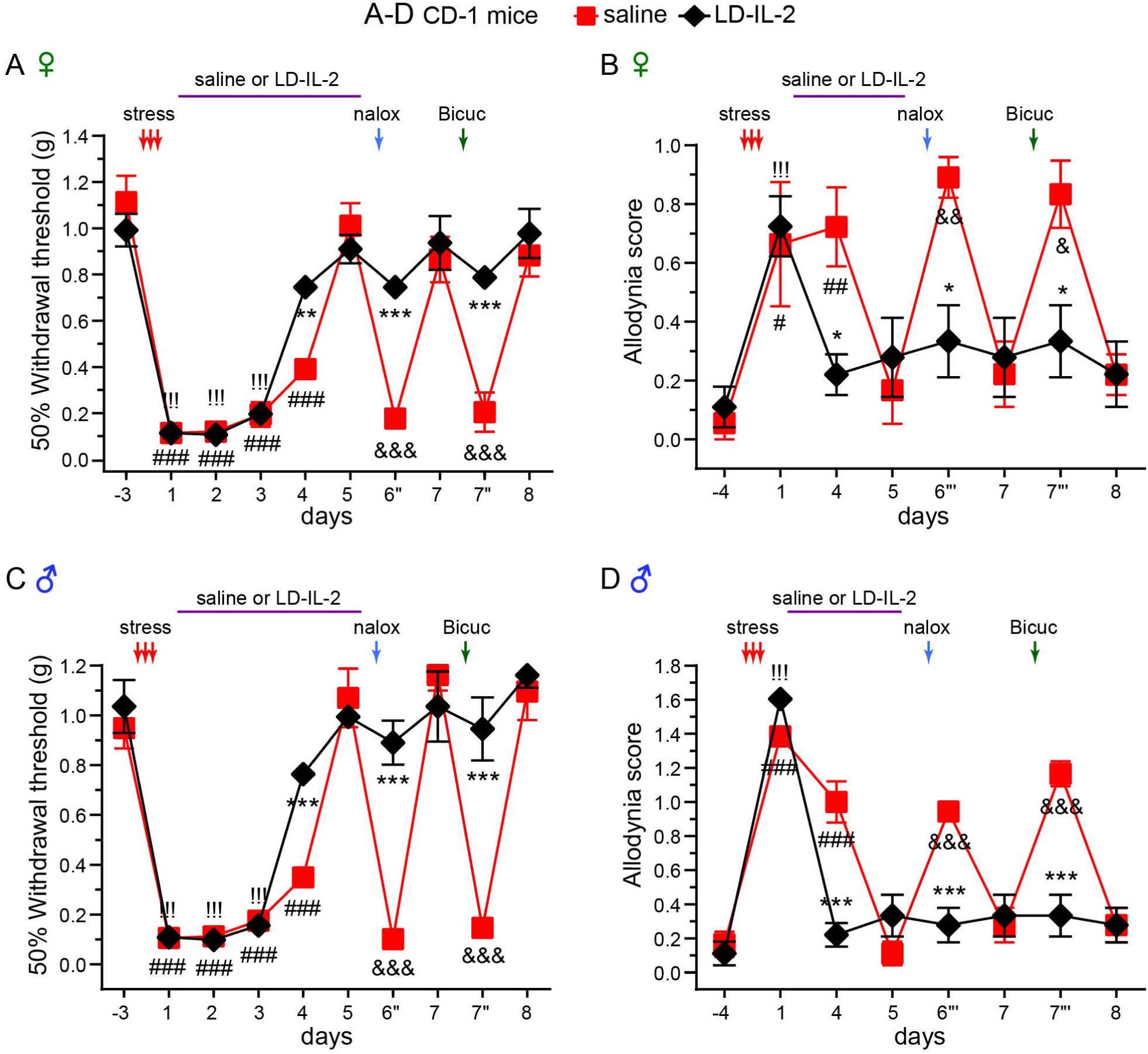
LD-IL-2 treatment prevents repetitive stress-induced latent sensitization on hindpaw skin. **(A-D)** Post-stress LD-IL-2 treatment prevented opioid and GABA_A_ receptor antagonists from reinstating the hypersensitivity to punctate **(A, C)** and dynamic **(B, D)** mechanical stimuli on hindpaw skin in female **(A-B)** and male **(C-D)** CD-1 mice (*n* = 6/group). \**P* < 0.05, \*\**P* < 0.01, \*\*\**P* < 0.001, saline versus LD-IL-2 groups; ^#^*P* < 0.05, ^##^*P* < 0.01, ^###^*P* < 0.001, compared with the pre-stress baseline thresholds in saline groups; ^!!!^*P* < 0.001, compared with the pre-stress baseline thresholds in LD-IL-2 group; ^&^*P* < 0.05, ^&&^*P* < 0.01, ^&&&^*P* < 0.001, compared with the values before each naloxone or Bicuc injection in the saline groups.

### 3.5. LD-IL-2 treatment prevents headache-related behavioral sensitization in stressed male and female mice

We went on to assess whether LD-IL-2 treatment is effective for stress-induced headache. First, female CD-1 mice received daily LD-IL-2 after repetitive stress. The resolution of acute sensitization was significantly accelerated after 4 days of treatment (Fig. 9A, day 5-9), when LD-IL-2-induced Treg expansion reached plateau [16; 91]. After the facial mechanical thresholds returned to baseline in both groups, neither SNP injection nor BLS exposure could re-establish facial skin hypersensitivity in LD-IL-2-treated mice, indicating that LD-IL-2 prevents the development of hyperalgesic priming. Likewise, administration of MNB, Bicuc or Sac did not reduce facial mechanical thresholds in the LD-IL-2 group (Fig. 9B), suggesting that peripheral opioid, GABA_A_ or GABA_B_ receptors are no longer required to maintain the pain-free state. These findings support the effectiveness of post-stress LD-IL-2 treatment in preventing the transition from acute to chronic headache.

**Figure 9.**
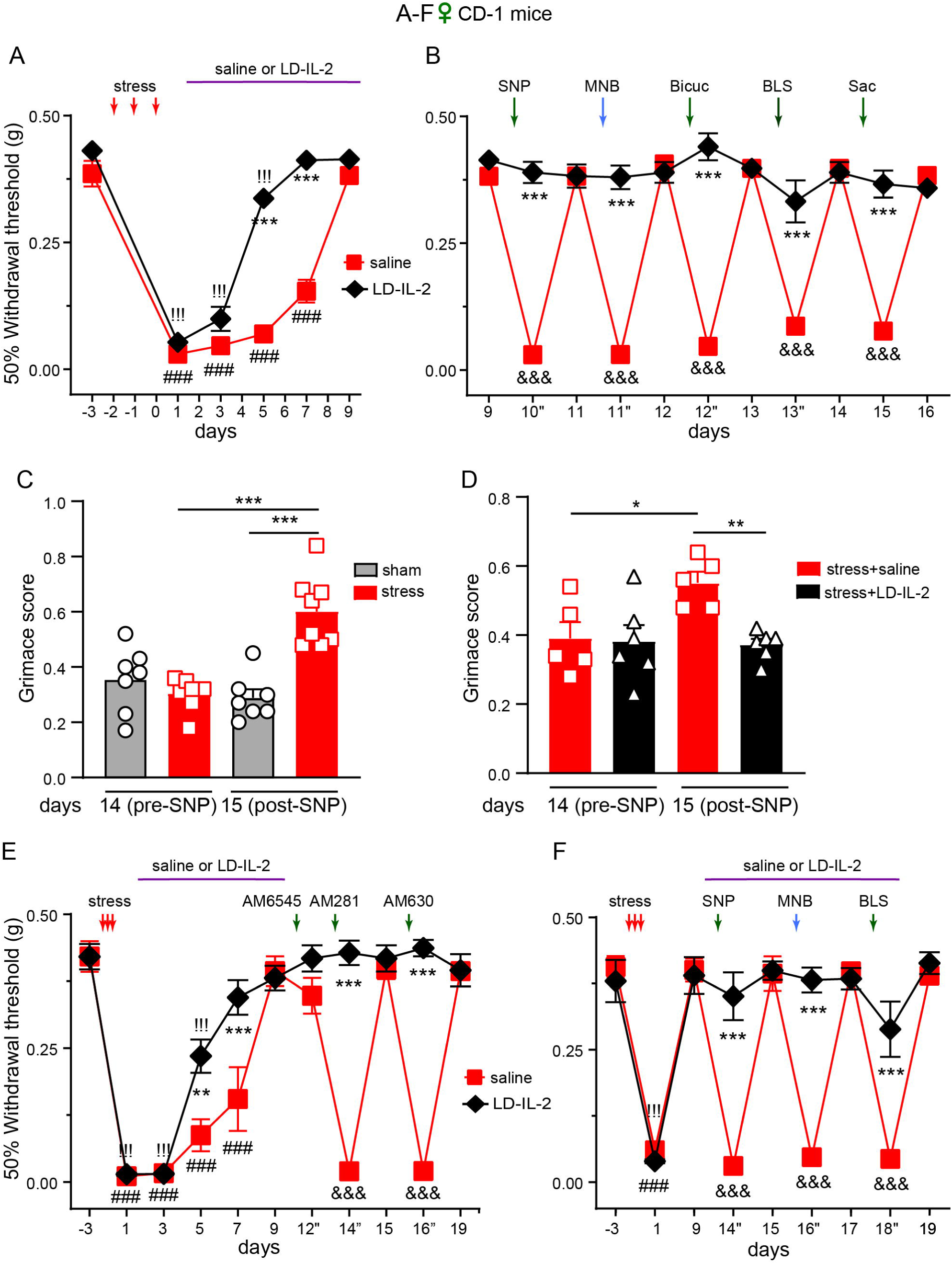
LD-IL-2 treatment prevents the development of headache-related chronic sensitization in stressed female mice. **(A)** LD-IL-2 accelerated the resolution of repetitive stress-induced acute sensitization in female CD-1 mice (*n* = 10/group). \*\*\**P* < 0.001, saline versus LD-IL-2 groups; ^###^*P* < 0.001, ^!!!^*P* < 0.001, compared with the day -3 baseline thresholds in saline and LD-IL-2 groups, respectively. **(B)** LD-IL-2 treatment prevented the development of stress-induced chronic sensitization in female CD-1 mice (same as in **A**). \*\*\**P* < 0.001, saline versus LD-IL-2 groups; ^&&&^*P* < 0.001, relative to the thresholds before SNP, MNB, Bicuc, BLS or Sac exposure in the saline group. **(C)** SNP administration increased facial grimace score in stressed female CD-1 mice but not in sham control (*n* = 7-8/group). \*\*\**P* < 0.001. **(D)** LD-IL-2 treatment prevented SNP-induced increase in facial grimace score in stressed female CD-1 mice (*n* = 5-6/group). \**P* < 0.05, \*\**P* < 0.01. **(E)** CB_1_R and CB_2_R antagonists could not evoke facial skin hypersensitivity in stressed mice that received LD-IL-2 treatment (*n* = 8 CD-1 females/group). \*\**P* < 0.01, \*\*\**P* < 0.001, saline versus LD-IL-2 groups; ^###^*P* < 0.001, ^!!!^*P* < 0.001, compared with the day -3 baseline thresholds in saline and LD-IL-2 groups, respectively; ^&&&^*P* < 0.001, relative to the day 9 thresholds before AM281 and day 15 thresholds before AM630 administration in the saline group. **(F)** Delayed LD-IL-2 treatment abolished SNP-, MNB-or BLS-evoked facial skin hypersensitivity in stressed mice (*n* = 10 CD-1 females/group). \*\*\**P* < 0.001, saline versus LD-IL-2 groups; ^###^*P* < 0.001, ^!!!^*P* < 0.001, compared with the day -3 baseline thresholds in saline and LD-IL-2 groups, respectively; ^&&&^*P* < 0.001, relative to the thresholds before SNP, MNB or BLS challenges in the saline group.

Next, we used the facial grimace score as a measure of nonreflexive, ongoing headache-related behaviors in female CD-1 mice (Fig. 9C-D) [4; 16; 44]. SNP had no effect on sham mice but significantly increased the grimace score in stressed mice (Fig. 9C), indicating that the subthreshold trigger elicits ongoing headache-related behaviors. In the second experiment, mice received daily LD-IL-2 treatment between day 1 and 9 post-stress. SNP increased the grimace score in saline-treated mice but had no effect on LD-IL-2-treated mice (Fig. 9D), suggesting that LD-IL-2 alleviates stress-induced ongoing headache.

We also found that the brain-penetrating CB_1_ receptor antagonist AM281 (2 mg/kg, i.p.) re-established facial skin hypersensitivity in stressed mice after the resolution of acute sensitization but the peripherally restricted CB_1_ receptor antagonist AM6545 (2 mg/kg, i.p.) had no effect (Fig. 9E, saline group, day 9-15), indicating that repetitive stress engages central but not peripheral CB_1_ receptor signaling to mask the latent sensitization. The CB_2_ receptor antagonist AM630 (2 mg/kg, i.p.) also elicited robust facial skin hypersensitivity in stressed mice (Fig. 9E, saline group, day 16”). Since AM630 is brain-penetrating, it is not possible to distinguish whether stress activates peripheral and/or central CB_2_ receptors. After LD-IL-2 treatment, stressed mice did not respond to either AM281 or AM630 (Fig. 9E, day 9-19, LD-IL-2 group), suggesting that LD-IL-2 prevents the persistent CB_1_ and CB_2_ receptor signaling in stressed mice.

To determine whether LD-IL-2 treatment is still effective after the chronic sensitization is fully established, we started the 9-day LD-IL-2 treatment after the resolution of acute sensitization. In saline-treated mice, SNP, BLS and MNB all evoked robust facial skin hypersensitivity, confirming the establishment of stress-induced chronic sensitization (Fig. 9F). Conversely, LD-IL-2-treated mice were fully protected from all these headache triggers (Fig. 9F). The fact that LD-IL-2 remained effective for well-established chronic sensitization highlights its therapeutic potential for stress-induced headache.

We also assessed the effects of LD-IL-2 in stressed male CD-1 mice. As in female mice, LD-IL-2-treated males exhibited faster recovery of acute sensitization (Fig. 10A) and were resilient to SNP, MNB or BLS challenges, indicating that LD-IL-2 treatment prevents the development of chronic sensitization in both sexes (Fig. 10B). When measuring the aversiveness/unpleasantness of headache, we found that all mice showed similar increase in facial pain score one day after repetitive stress, before the initiation of LD-IL-2 treatment (Fig. 10C, day 1). After 9 days of saline or LD-IL-2 treatment, SNP administration increased facial pain score in saline-treated mice but not in the LD-IL-2 group (Fig. 10D, day 12), suggesting that LD-IL-2 attenuates the unpleasantness of headache. Collectively, our findings support LD-IL-2 as a potential disease-modifying treatment for stress-induced persistent headache and widespread pain in both males and females.

**Figure 10.**
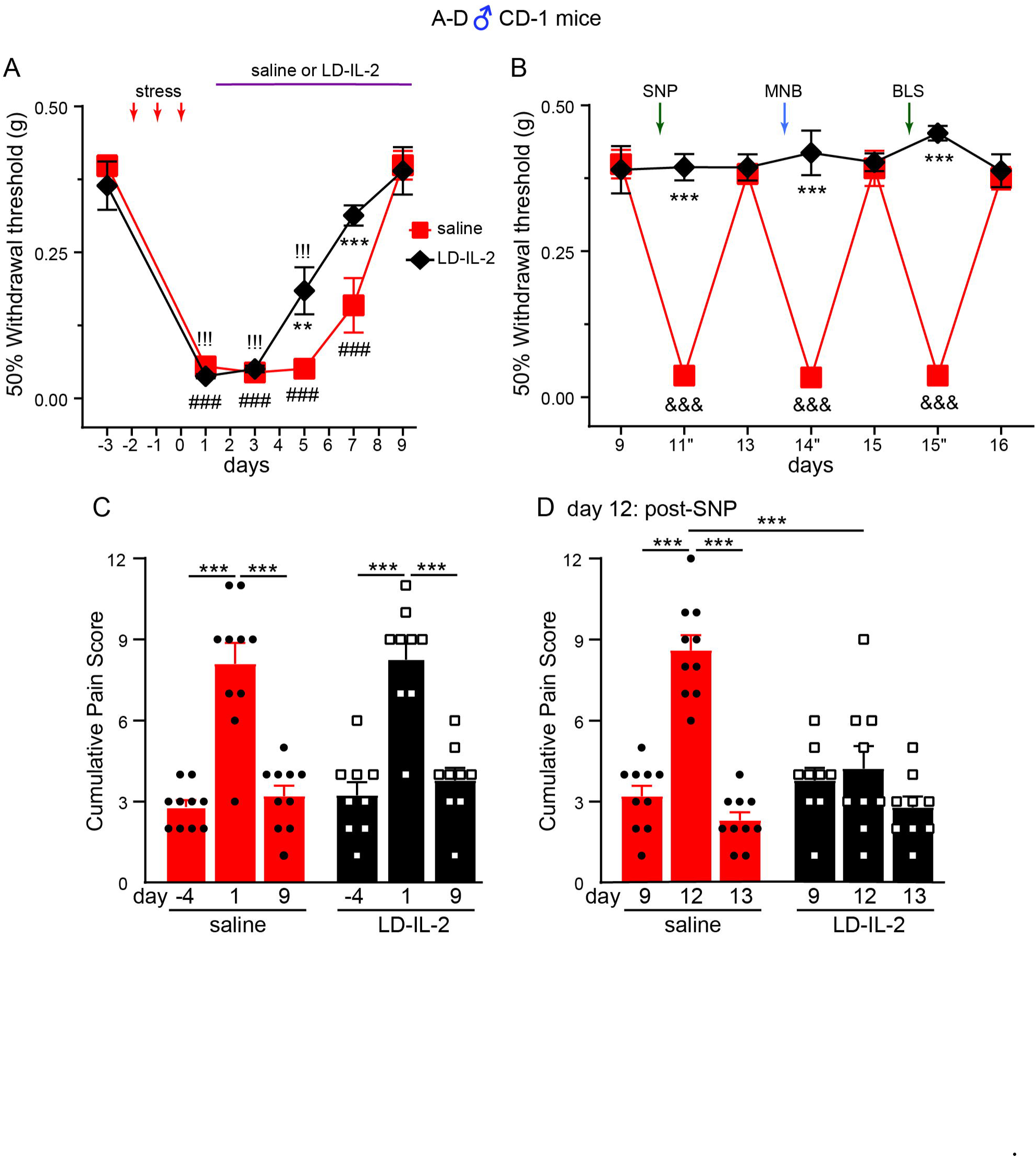
LD-IL-2 treatment exhibits similar therapeutic effects in stressed male mice. **(A)** LD-IL-2 accelerated the resolution of stress-induced acute sensitization in male CD-1 mice (*n* = 9-10/group). \*\*\**P* < 0.001, saline versus LD-IL-2 groups; ^###^*P* < 0.001, ^!!!^*P* < 0.001, compared with the day -3 baseline thresholds in saline and LD-IL-2 groups, respectively. **(B)** LD-IL-2 treatment prevented the development of stress-induced chronic sensitization in male CD-1 mice (same as in **A**). \*\*\**P* < 0.001, saline versus LD-IL-2 groups; ^&&&^*P* < 0.001, relative to the thresholds before SNP, MNB, or BLS exposure in the saline group. **(C)** Mice exhibited similar stress-induced increase in facial pain score prior to LD-IL-2 treatment (same mice as in **A**).\*\*\**P* < 0.001. **(D)** LD-IL-2 treatment prevented SNP-induced increase in facial pain score in stressed male CD-1 mice (same as in **A**). Mice received a 2^nd^ dose of SNP (0.3 mg/kg, i.p.) on day 12. Facial pain scores were quantified 3 hours post-SNP. \*\*\**P* < 0.001.

### 3.6. LD-IL-2/Treg engages multiple peripheral cytokine signaling pathways but does not depend on opioid receptor activation

We used DEREG mice to test whether LD-IL-2 inhibits stress-induced behaviors through endogenous Treg cells. After repetitive stress, male and female DEREG mice received two rounds of saline or DT administration during the daily LD-IL-2 treatment session. In saline-treated mice with intact Treg cells, LD-IL-2 accelerated the resolution of acute facial mechanical hypersensitivity (Fig. 11A) as in CD-1 mice (Fig. 9A and 10A). To test whether LD-IL-2 treatment protects mice against stronger pain triggers, we treated mice with higher doses of SNP (1 mg/kg) and PGE_2_ (300 ng). In mice with intact Treg cells, LD-IL-2 treatment prevented these stimuli from re-establishing facial or hindpaw sensitization (Fig. 11A-B). Conversely, the effects of LD-IL-2 were abolished in DT-treated, Treg-depleted mice. Stress-induced acute facial skin hypersensitivity was significantly prolonged (Fig. 11A, day -3 ∼ 16), like Treg-depleted mice without LD-IL-2 treatment (Fig. 6A). SNP and PGE_2_ evoked robust facial and hindpaw hypersensitivity (Fig. 11A-B). These results confirm that endogenous Treg cells mediate the therapeutic effects of LD-IL-2 after repetitive stress.

**Figure 11.**
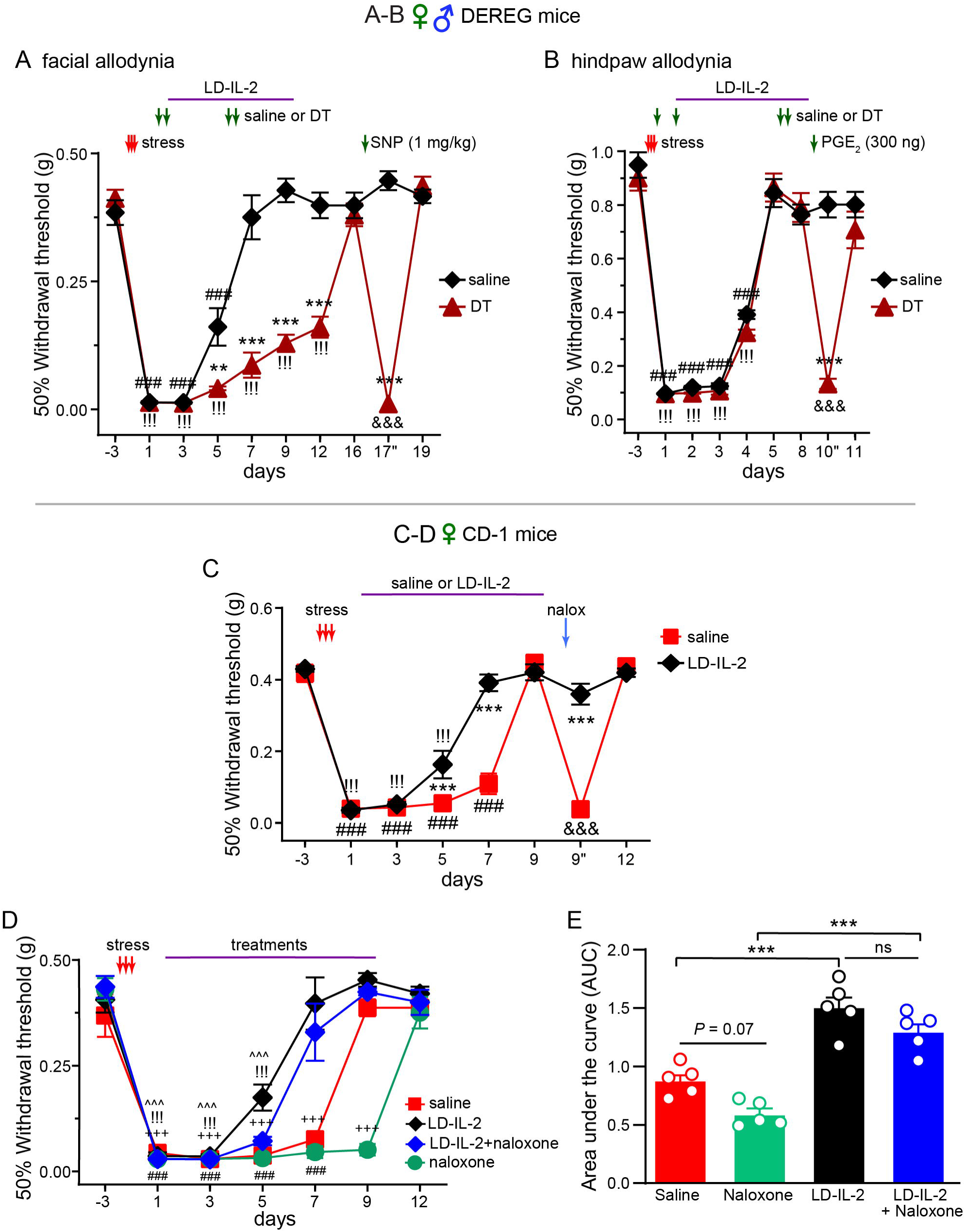
Treg cells mediate the effects of LD-IL-2 independent of endogenous opioid receptor signaling. **(A-B)** Depletion of Treg cells through repeated DT administration abolished the effects of LD-IL-2 on stress-induced facial and hindpaw mechanical hypersensitivity in DEREG mice (*n* = 4 male and 3-4 female mice/group). Note that higher doses of SNP (1 mg/kg) and PGE_2_ (300 ng) were administrated during the chronic phase to test whether LD-IL-2 was effective in preventing re-sensitization by stronger stimuli. \*\*\**P* < 0.001, saline versus DT; ^###^*P* < 0.001, ^!!!^*P* < 0.001, compared with the day -3 baseline thresholds within the saline and DT groups, respectively; ^&&&^*P* < 0.001 within the DT group, compared with the day 16 thresholds before SNP injection in **A** and compared with the day 8 thresholds before PGE_2_ injection in **B**. **(C)** Naloxone-induced facial skin hypersensitivity was abolished in stressed female CD-1 mice that received LD-IL-2 treatment (*n* = 8/group). \*\*\**P* < 0.001, saline versus LD-IL-2 groups; ^###^*P* < 0.001, ^!!!^*P* < 0.001, compared with the day -3 baseline thresholds in saline and LD-IL-2 groups, respectively; ^&&&^*P* < 0.001, relative to the thresholds before naloxone administration in the saline group. **(D)** Naloxone administration (2 mg/kg/day for 9 days, i.p.) delayed the resolution of stress-induced acute sensitization in male CD-1 mice (*n* = 5/group). However, co-administration of naloxone did not prevent LD-IL-2 from facilitating the resolution of acute sensitization. ^###^*P* < 0.001, ^!!!^*P* < 0.001, ^+++^*P* < 0.001, ^^^^^*P* < 0.001, compared with the day -3 baseline thresholds in saline, LD-IL-2, naloxone, and LD-IL-2+naloxone groups, respectively. **(E)** Comparing the areas under the curve (AUC, day -3 to 12) of individual mice in **(D)** confirmed that co-administration of naloxone did not prevent LD-IL-2 from facilitating the resolution of acute sensitization. \*\*\**P* < 0.001.

We went on to investigate LD-IL-2’s molecular MoA. Two recent studies report that enkephalin-producing Treg cells inhibit nociception through endogenous opioid receptor signaling [58; 59]. However, LD-IL-2 treatment prevented MNB or naloxone from evoking facial and hindpaw mechanical hypersensitivity in stressed mice (Fig. 8A-D, 9B, 9F, 10B, and Fig. 11C), indicating that neither peripheral nor central opioid receptor signaling is required to maintain normal nociceptive responses in LD-IL-2-treated mice. To address whether LD-IL-2 facilitates the resolution of stress-induced acute sensitization through opioid receptor signaling, mice received naloxone daily to continuously block all central and peripheral opioid receptor subtypes during LD-IL-2 treatment (Fig. 11D, day 1-9). Naloxone administration alone prolonged stress-induced sensitization (Fig. 11D, day 9, saline versus naloxone), indicating that endogenous opioid receptors are activated during the acute phase to accelerate its resolution. Importantly, in mice that received both LD-IL-2 and naloxone, the resolution of acute sensitization followed the similar time course as those received LD-IL-2 alone, significantly faster than the saline or naloxone group (Fig. 11D day 7-9, and 11E). We conclude that the beneficial effects of LD-IL-2 on stress-induced acute and chronic sensitization are not dependent on opioid receptor signaling.

One of the main mechanisms that Treg cells use for immunosuppression is through the secretion of cytokines including TGF-β1 and IL-10 [20]. In a mouse model of nitric oxide-induced chronic migraine, we have shown that LD-IL-2 reverses headache-related sensitization through peripheral IL-10 and TGF-β1 pathways [33]. Although IFN-γ is generally considered a pro-inflammatory cytokine, Treg-produced IFN-γ has been shown to exert stable antiviral and immunosuppressive functions [17; 29; 74]. To investigate whether these cytokines mediate the therapeutic effects of LD-IL-2 in stressed mice, we treated female CD-1 mice with neutralizing antibodies against IL-10, IFN-γ, or against all TGF-β isoforms. Baseline mechanical responses were not altered by either antibody (Fig. 12A-C, day -5 versus -3). Stress-induced acute or chronic sensitization were not changed by the anti-IFN-γ or anti-IL-10 antibodies (Fig. 12A-B) but were drastically prolonged by the anti-TGF-β antibody (Fig. 12C), indicating that the peripheral TGF-β signaling pathway facilitates the resolution of stress-induced sensitization.

**Figure 12.**
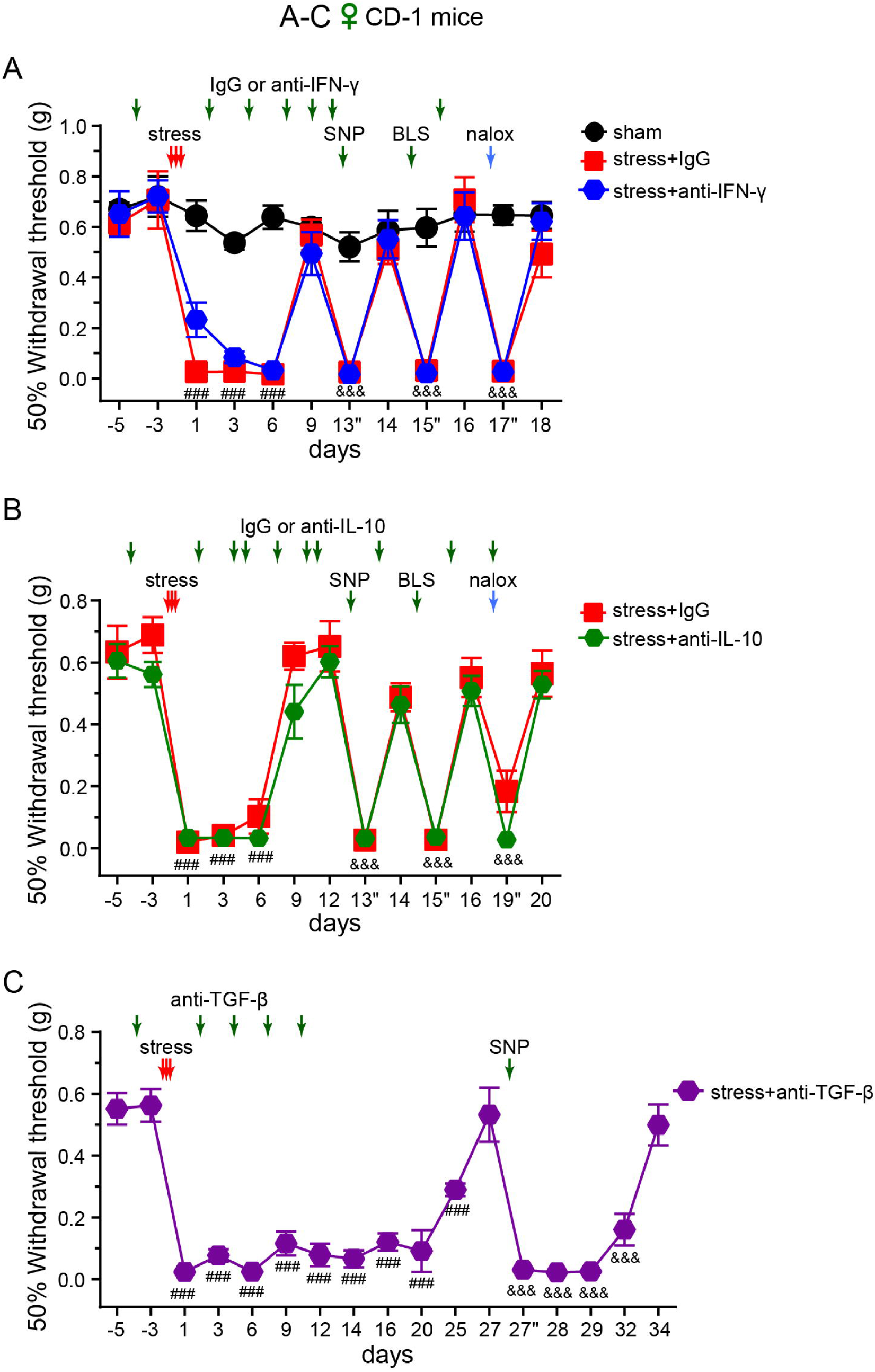
Repetitive stress recruits peripheral TGF-β signaling pathway to facilitate the resolution of headache-related sensitization. **(A)** The neutralizing antibody against IFN-γ (anti-IFN-γ) did not alter baseline mechanical responses or stress-induced behaviors in female CD-1 mice (*n* = 6/group). Anti-IFN-γ and control IgGs were i.p. administered (200 µg/mouse) every 3 days. ^###^*P* < 0.001, compared with the day -5 baseline thresholds within the stress+IgG and stress+anti-IFN-γ groups. ^&&&^*P* < 0.001, compared with the values before SNP, BLS or naloxone challenges within the stress+IgG and stress+anti-IFN-γ groups. **(B)** Neither baseline mechanical responses nor stress-induced behaviors in female CD-1 mice were affected by the neutralizing antibody against IL-10 (anti-IL-10, *n* = 4 for stress+IgG and 8 for stress+anti-IL-10 groups. Anti-IL-10 and control IgGs were i.p. administered (200 µg/mouse) every 2 days. ^###^*P* < 0.001, compared with the day -5 baseline thresholds within the stress+IgG and stress+anti-IL-10 groups. ^&&&^*P* < 0.001, compared with the values before SNP, BLS or naloxone challenges within the stress+IgG and stress+anti-IL-10 groups. **(C)** The neutralizing antibody against all TGF-β isoforms (anti-TGF-β) did not affect baseline mechanical withdrawal thresholds, but prolonged stress-induced acute and chronic sensitization in female CD-1 mice (*n* = 7). Note that mice in **B** and **C** were assessed in parallel but plotted separately for clarify. ^###^*P* < 0.001, ^&&&^*P* < 0.001, compared with the day-5 baseline thresholds and day 27 thresholds prior to SNP injection, respectively.

Interestingly, co-administration of either neutralizing antibody during LD-IL-2 treatment abolished LD-IL-2’s effects on stress-induced chronic sensitization (Fig. 13A-B). In mice that received anti-IFN-γ or anti-IL-10 antibody, SNP, BLS and naloxone evoked robust facial skin hypersensitivity (Fig. 13A), like those in stressed mice without LD-IL-2 treatment (Fig. 12A). In mice that received the anti-TGF-β antibody, SNP-induced sensitization did not fully resolve till day 21 (Fig. 13B). Subsequent BLS exposure had no effect, likely because stress-induced hyperalgesic priming is no longer present at this point (Fig. 1B). Thus, LD-IL-2-treated mice engage at least three peripheral cytokine signaling pathways – IFN-γ, IL-10 and TGF-β – to prevent stress-induced chronic sensitization.

**Figure 13.**
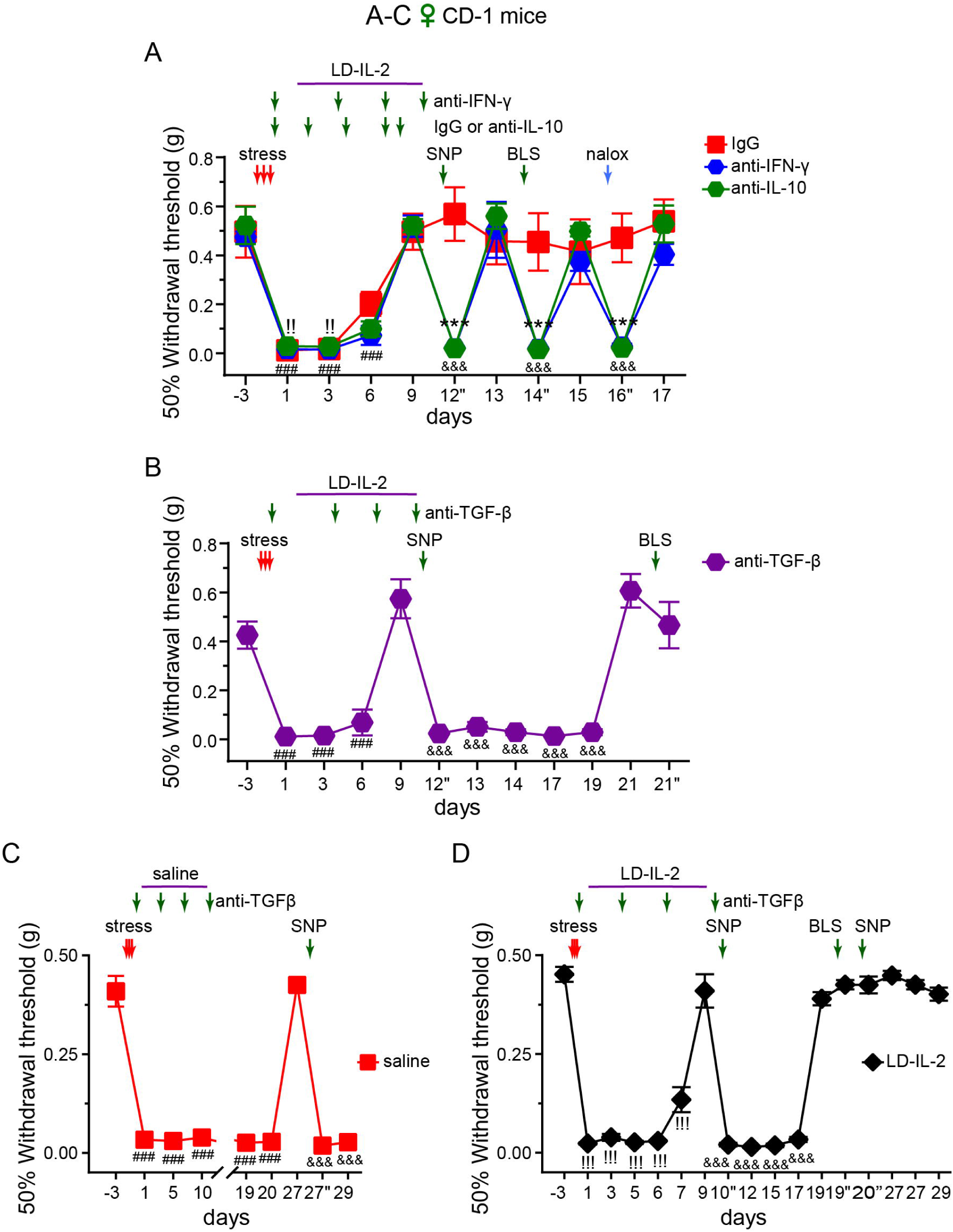
LD-IL-2 prevents stress-induced chronic sensitization through peripheral IFN-γ, IL-10 and TGF-β signaling pathways. **(A)** Neutralizing either IFN-γ or IL-10 negated the effects of LD-IL-2 on stress-induced chronic sensitization (*n* = 4-5/group). \*\*\**P* < 0.001, stress+IgG versus stress+anti-IL-10 and stress+anti-IFN-γ groups; ^###^*P* < 0.001, compared with the day -3 baseline thresholds within the stress+anti-IL-10 and stress+anti-IFN-γ groups; ^!!^*P* < 0.01, compared with the day -3 baseline thresholds in the stress+IgG group; ^&&&^*P* < 0.001, compared with the values before SNP, BLS or naloxone challenges within the stress+anti-IFN-γ and stress+anti-IL-10 groups. **(B)** Neutralizing TGF-β abolished the effect of LD-IL-2 on stress-induced chronic sensitization, prolonging SNP-induced facial skin hypersensitivity (*n* = 5). Note that mice in **A** and **B** were assessed in parallel but plotted separately for clarity. ^###^*P* < 0.001, ^&&&^*P* < 0.001, compared with the day -3 and day 9 thresholds, respectively. **(C-D)** The effects of anti-TGF-β neutralizing antibody were independently verified by another researcher (*n* = 7/ female CD-1 group). Note that mice in **C** and **D** were tested in parallel but plotted separately for clarity. ^###^*P* < 0.001, ^!!!^*P* < 0.001, compared with the day -3 baseline thresholds within saline and LD-IL-2 groups; ^&&&^*P* < 0.001, compared with the day 27 thresholds in **A** or day 9 thresholds in **B**.

Another researcher independently verified the effects of blocking peripheral TGF-β signaling (Fig. 13C-D). Unlike the experiment in Fig. 12C (day -5), all mice received anti-TGF-β antibody after repetitive stress. This was sufficient to delay the resolution of headache-related sensitization (Fig. 13C). Notably, in LD-IL-2-treated mice, anti-TGF-β antibody prolonged the duration of SNP-induced facial skin hypersensitivity but not the acute sensitization (Fig. 13B and 13D versus the IgG group in Fig. 13A). These results suggest that, in the absence of TGF-β signaling, LD-IL-2 can employ other mechanisms (e.g. IFN-γ and/or IL-10 pathways) to resolve acute sensitization. During the chronic phase, all three cytokine pathways are required to block trigger-induced re-sensitization. Nonetheless, only peripheral TGF-β signaling is indispensable for limiting the response duration to subthreshold stimuli. Lastly, mice in Fig. S5B did not respond to the 2^nd^ SNP injection on day 20 post-stress, suggesting that loss of peripheral TGF-β signaling does not prolong the duration of hyperalgesic priming.

### 3.7. LD-IL-2 is more effective than anti-CGRP treatment in preventing repetitive stress-induced chronic sensitization

Drugs targeting CGRP or its receptor have shown promising anti-migraine effects in both randomized clinical trials and real-world studies [24; 64; 71]. This prompted us to compare the effectiveness of LD-IL-2 with a chimeric neutralizing antibody against CGRP [72]. In a previous study, one injection of an anti-CGRP antibody at 30 mg/kg blocks acute post-traumatic headache-related behaviors for 6 days [62]. We therefore treated mice with i.p. injection of the chimeric anti-CGRP antibody (30 mg/kg) on day 10 post-stress and tested their responses to three subthreshold triggers within 5 days (Fig. 14A, day 11”-15”). SNP became ineffective but Bicuc and BLS could still induce re-sensitization after anti-CGRP treatment (Fig. 14A). This is consistent with previous reports that anti-CGRP antibody blocks SNP-induced sensitization after repetitive stress but is ineffective for BLS-induced post-traumatic headache [4; 62]. Conversely, LD-IL-2 treatment abolished the effects of all three triggers (Fig. 14A), as in the previous experiment (Fig. 9B). The difference between anti-CGRP antibody and LD-IL-2 unlikely results from the faster resolution of acute sensitization by LD-IL-2, as delaying LD-IL-2 session to day 9-17 post-stress still prevents BLS-induced facial skin hypersensitivity (Fig. 9F). We therefore conclude that LD-IL-2 is more effective than anti-CGRP treatment in preventing repetitive stress-induced chronic sensitization.

**Figure 14.**
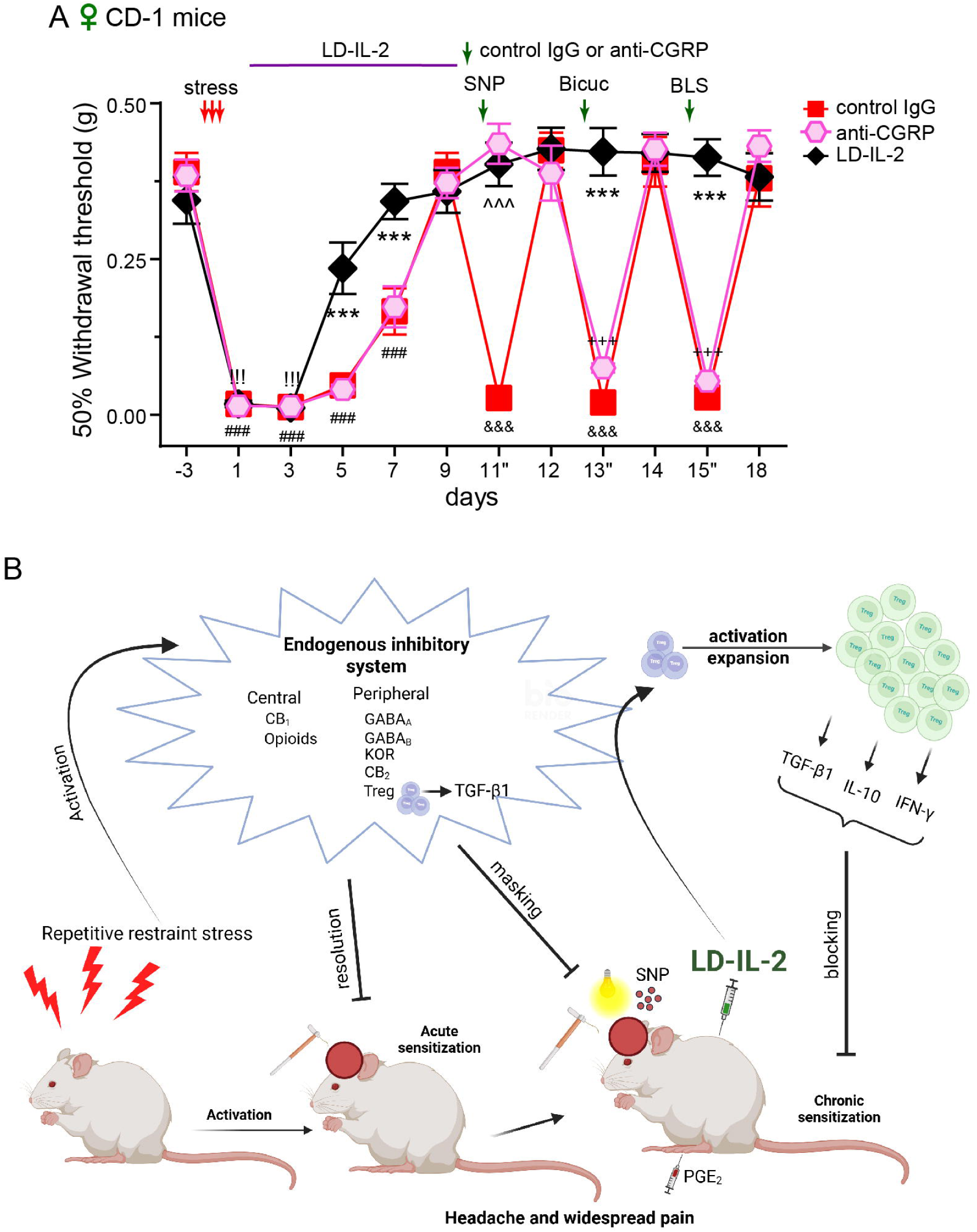
LD-IL-2 is more effective than anti-CGRP treatment in preventing repetitive stress-induced chronic sensitization. **(A)** Bicuc-and BLS-induced facial skin hypersensitivity during the chronic sensitization phase was prevented by LD-IL-2 treatment but not by the chimeric anti-CGRP antibody in female CD-1 mice (*n* = -7/group). \*\*\**P* < 0.001, LD-IL-2 versus anti-CGRP and control IgG; ^^^^^*P* < 0.001, LD-IL-2 versus control IgG only; ^###^*P* < 0.001, compared with the day -3 baseline thresholds in control IgG and anti-CGRP groups; ^!!!^*P* < 0.001, compared with the day -3 baseline thresholds in the LD-IL-2 group; ^&&&^*P* < 0.001, ^+++^*P* < 0.001, compared with the values before SNP, Bicuc, or BLS challenges within the control IgG and anti-CGRP groups, respectively. **(B)** Graphic summary of the main results from this study (created using BioRender).

## 4. Discussion

Despite the well-established link between stress and comorbid migraine and widespread somatic pain [94], the endogenous protective mechanisms that actively suppress pain and maintain interictal stability have not been systematically explored. The present study addressed this knowledge gap (Fig. 14B). We confirmed that repetitive restraint stress in mice produced robust acute facial and hindpaw mechanical hypersensitivity mechanistically related to migraine headache and widespread somatic pain, followed by a hyperalgesic priming period during which subthreshold triggers (SNP, BLS and PGE_2_) re-established cutaneous allodynia. In the absence of triggering events, hyperexcitation of the head and body pain circuits was masked by concurrent peripheral opioid and GABAergic signaling across sexes and strains, underscoring its broad biological relevance of the latent sensitization state.

### 4.1. Common versus selective inhibitory mechanisms for headache and body pain

Parallel assessment of cephalic and somatic allodynia revealed that the same repetitive stress paradigm induced longer-lasting sensitization in the trigeminovascular pathway than in somatic pain circuits, providing a wider time window to identify peripheral inhibitory pathways masking the latent cephalic sensitization. Our findings identify a previously unrecognized division of labor among endogenous inhibitory systems following repetitive stress. Within the first 3 weeks after stress, peripheral GABA_A_ and GABA_B_ receptor signaling suppresses latent sensitization, whereas endocannabinoids act through central but not peripheral CB_1_ receptors. Additionally, repetitive stress activated peripheral KOR signaling to suppress chronic cephalic sensitization. Results from this work and a previous study using the same model [56] implicate peripheral DOR and KOR, but not MOR, in inhibiting chronic headache-related latent sensitization. Beyond three weeks, central opioid signaling takes over to continuously mask the hyperexcitation of pain circuits. Future studies should determine the central opioid receptor subtypes involved, whether CB_2_ receptors are activated centrally or peripherally, and whether central GABA_A_ and GABA_B_ receptors are activated alongside opioid receptors.

Consistent with our prior finding that Treg cells facilitate recovery from post-traumatic headache, systemic Treg depletion markedly delayed the resolution of stress-induced acute facial skin hypersensitivity and prolonged SNP-evoked re-sensitization. In contrast, Treg depletion had no effect on hindpaw allodynia, revealing a previously unrecognized selectivity of Treg-mediated modulation of trigeminal pain after repetitive stress. Even with additional inhibition from Treg cells, facial sensitization persisted longer than hindpaw allodynia, underscoring the vulnerability of headache circuits to repetitive stress. Our findings suggest that maintaining the pain-free interictal state relies on the coordinated activity of Treg cells together with peripheral KOR, GABA_A_, GABA_B_, central CB_1_, CB_2_, and central opioid receptor signaling at later stage. Compromising any single component unmasks latent sensitization and increase risk of migraine attacks. Whether impairment of endogenous inhibition systems underlies the pathophysiology of drug-resistant and refractory migraine, rendering patients unresponsive to single-pathway targeting drugs [70], warrants further investigation.

Despite the engagement of multiple inhibitory systems, stressed mice remained responsive to subthreshold stimuli, suggesting that therapeutic enhancement of inhibition beyond basal levels may be necessary to “reset” pain circuits. Given that frequent use of brain-penetrating opioids and barbiturates increases the risk of medication overuse headache [2; 93], peripherally restricted agonists or positive allosteric modulators for KOR, GABA_A_, GABA_B_, and cannabinoid receptors merit evaluation as migraine prophylactic treatments [19; 68]. The peripherally-acting KOR agonist difelikefalin is already approved for treating moderate-to-severe pruritus in humans. Since it cross-reacts with mouse KOR, future preclinical studies will assess whether it can be repurposed as chronic headache treatment.

### 4.2. Therapeutic efficacy and flexible time window of LD-IL-2

In our previous work, systemic LD-IL-2 treatment robustly expands Treg cells in dura, trigeminal ganglion, sciatic nerves and DRG tissues relevant to headache and somatic pain [16; 36; 73; 91]. In mice with established stress-induced hindpaw hypersensitivity, LD-IL-2 treatment accelerated its resolution and prevented chronic sensitization in both sexes. Likewise, post-stress LD-IL-2 facilitated the resolution of acute facial skin hypersensitivity and prevented migraine triggers SNP and BLS from re-establishing sensitization. After LD-IL-2, blocking other inhibitory pathways – including GABA_A_, GABA_B_, CB_1_, CB_2_ and opioid receptors – no longer reinstated facial skin hypersensitivity, suggesting a disease-modifying effect of LD-IL-2. LD-IL-2 also normalized facial pain score and grimace score in stressed mice, suggesting mitigation of both sensory and affective components of ongoing headache [4; 8; 14; 16; 44; 72]. Moreover, delayed LD-IL-2 treatment was still effective after the establishment of stress-induced chronic sensitization (Fig. 9F), suggesting a flexible therapeutic time window. These findings support LD-IL-2 and other Treg-enhancing strategies as potential disease-modifying treatments for stress-induced migraine and comorbid body pain beyond symptomatic management, an exciting possibility requiring further validation.

### 4.3. LD-IL-2 operates independently of canonical inhibitory pathways

We found that Treg cells mediate the therapeutic benefits of LD-IL-2 on stress-induced sensitization, consistent with its cellular mechanism in other disease models [16; 41; 76; 91]. Regarding molecular mechanisms, both enkephalin-DOR signaling and cytokine TGF-β, IL-10, and interleukin-32 pathways have been shown to mediate LD-IL-2/Treg’s analgesic effects [23; 33; 58; 59]. Although blocking both central and peripheral opioid signaling with naloxone prolonged stress-induced acute sensitization, coadministration of naloxone during LD-IL-2 treatment still accelerated the resolution of acute sensitization (Fig. 11D-E). Furthermore, blocking opioid, GABA_A_, GABA_B_, CB_1_, or CB_2_ receptor signaling after LD-IL-2 failed to re-establish facial or hindpaw mechanical hypersensitivity in stressed mice (Fig. 9B, 9E and 11C). These results point to a molecular mechanism of LD-IL-2 independent of endogenous opioid, GABA and cannabinoid signaling pathways for stress-induced pain.

Stressed mice without LD-IL-2 treatment primarily relied on peripheral TGF-β, but not IL-10 or IFN-γ, signaling to facilitate the resolution of cephalic allodynia. In contrast, LD-IL-2 acted through Treg cells to recruit all three cytokines to reverse stress-induced chronic sensitization. Neutralizing any single cytokine abolished LD-IL-2’s protective effects, permitting subthreshold stimuli to reinstate facial hypersensitivity. Specifically, while all three cytokines cooperatively regulate the response threshold to headache triggers, TGF-β signaling alone controlled response duration during the chronic phase. Conversely, neutralizing TGF-β did not prolong acute sensitization in LD-IL-2-treated mice, suggesting that enhanced IL-10 and IFN-γ signaling is sufficient to support its resolution. We therefore propose that LD-IL-2-expanded Treg cells concomitantly enhance peripheral TGF-β1, IL-10, and IFN-γ signaling to prevent stress-induced headache. Whether LD-IL-2/Treg unitizes the same mechanisms to control stress-induced somatic pain remains to be determined, but it likely is independent of opioid and GABAergic signaling (Fig. 8).

### 4.4. LD-IL-2 outperforms the chimeric anti-CGRP antibody

While drugs targeting the CGRP signaling pathway have revolutionized migraine treatment, 30-50% of patients do not achieve a clinically significant response [24; 64; 71]. Consistent with this limitation, the chimeric anti-CGRP antibody prevented SNP-induced re-sensitization in stressed mice but did not block Bicuc-or BLS-induced cephalic allodynia (Fig. 14A). In contrast, LD-IL-2 abolished re-sensitization across all tested triggers and replaced canonical inhibitory pathways in masking latent sensitization. This broader effect likely reflects LD-IL-2’s capacity to target Tregs and engage multiple cytokine pathways, thereby providing superior, disease-modifying protection compared to therapies targeting individual molecules.

### 4.5. Limitations of the study

Several limitations warrant future investigation. First, we did not identify the cellular sources of TGF-β1, IL-10, or IFN-γ. Follow-up studies should determine whether LD-IL-2 increases release of these cytokines directly from Treg cells or indirectly from other cells regulated by Tregs. Second, the downstream cellular targets of these cytokines remain uncertain. Chronic stress increases dural T cells in female rats and dural macrophages in male rats [55]. A recent study reports that macrophages contribute to migraine-related behavioral sensitization in stressed male and female mice [57]. Given Treg’s role as a master immune regulator, further studies will investigate whether LD-IL-2/Treg acts by restraining T cell or macrophage activity. Alternatively, cytokines may act directly on primary afferent neurons. Our prior work suggests that activation of TGF-β1 and IL-10 receptors in trigeminal ganglion neurons reverses chronic migraine-related sensitization in LD-IL-2-treated mice [33]. Although generally considered a pro-inflammatory cytokine, Treg cells can produce IFN-γ under certain disease states [17; 29; 74]. A recent study proposes that IFN-γ may reduce somatic pain by enhancing tetrodotoxin-sensitive Na^+^ currents in DRG neurons, causing membrane depolarization and inactivation of tetrodotoxin-insensitive Na^+^ channels, thereby eliminating interleukin-17A-induced hyperexcitation [61]. Future studies will determine whether similar neuronal mechanisms mediate the effects of LD-IL-2 on stress-induced headache and body pain. A third limitation is that only one stress model was used. It is important to examine whether our findings can be generalized to headache and widespread pain induced by other stressors in future studies.

## Conclusions

Taken together, the present study reveals multiple peripheral and central inhibitory pathways acting concertedly to mask repetitive stress-induced latent sensitization. Compromising any single pathway may increase migraine susceptibility. LD-IL-2 eliminates stress-induced latent sensitization through targeting Treg cells to recruit peripheral TGF-β1, IL-10, and IFN-γ signaling pathways. By bypassing central opioid, GABA and cannabinoid signaling, LD-IL-2 carries minimal risk of medication overuse headache. Together with its well-established safety profile in clinical trials for other diseases, these findings strongly support further translational evaluation of LD-IL-2 as a novel, disease-modifying treatment for stress-induced headache and widespread pain.

## Acknowledgements

The authors thank Dr. Jennifer Robblee for valuable discussions and Ms. Li Feng for excellent technical support.

## Data availability statement

All data are available upon reasonable request to the corresponding author.

## Figure legends

**Supplementary Figure 1.** Repetitive stress induces acute sensitization and hyperalgesic priming in male CD-1 mice (*n* = 6, same as the saline group in **Fig. 7C**). ^###^*P* < 0.001, compared with the day -3 baseline thresholds. Day 7”-9”, stress-induced hyperalgesic priming revealed by intraplantar injection of PGE_2_. ^&&&^*P* < 0.001, relative to the thresholds before each PGE_2_ injection.

**Supplementary Table 1.** Statistical analysis in individual experiments.

