## Supplementary figures and images for "Low-Dose Interleukin-2 Resolves Stress-Induced Chronic Sensitization Independent of Opioid Receptor Signaling"

### Supplementary Fig. 1

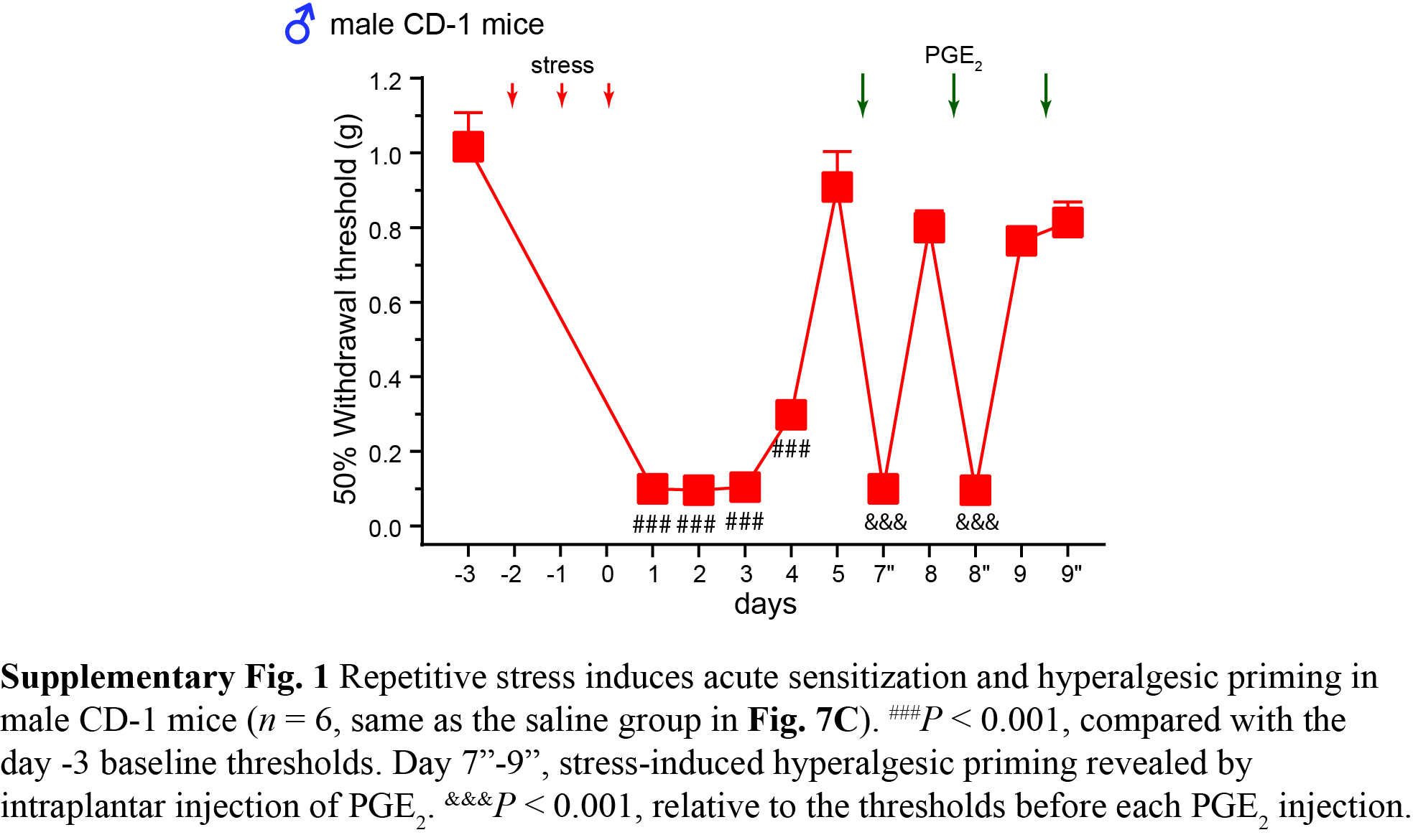
