## Supplementary Table 1 for "Low-Dose Interleukin-2 Resolves Stress-Induced Chronic Sensitization Independent of Opioid Receptor Signaling"

Supplementary Table 1. Statistical analysis in individual experiments.

| Figure | Statistical analysis | Post hoc analysis |
| --- | --- | --- |
| 1A | Two-way RM ANOVA, *P* < 0.001 for group (F [1, 10] = 214.3)  time (F [5, 50] = 111.2)  group x time interaction (F [5, 50] = 102.8)  One-way RM ANOVA:  Sham: *P* = 0.15, F [5, 25] = 1.774  Stress: *P* < 0.001, F [5, 25] = 142.1 | Bonferroni for multiple comparison:  ^***^*P* < 0.001, sham versus stress  ^###^*P* < 0.001, compared with the day -3 baseline threshold in the stress group |
| 1B | Two-way RM ANOVA, *P* < 0.001 for group (F [1, 10] = 59.66)  time (F [15, 150] = 38.14)  group x time interaction (F [15, 150] = 41.81)  One-way RM ANOVA:  sham: *P* = 0.93, F [15, 75] = 0.50  stress: *P* < 0.001, F [15, 75] = 54.63 | Bonferroni for multiple comparison:  ^***^*P* < 0.001, sham versus stress  ^&&&^*P* < 0.001, compared with the values before SNP, MNB, Bicuc, BLS and Sac challenges in the stress group |
| 1C | Two-way RM ANOVA, *P* < 0.001 for group (F [1, 10] = 142.2)  time (F [15, 150] = 24.48)  group x time interaction (F [15, 150] = 22.18)  One-way RM ANOVA:  sham: *P* = 0.02, F [15, 75] = 2.03  stress: *P* < 0.001, F [15, 75] = 41.80 | Bonferroni for multiple comparison:  ^***^*P* < 0.001, sham versus stress  ^###^*P* < 0.001, compared with the day -3 baseline threshold in the stress group  ^&&&^*P* < 0.001, compared with the values before naloxone, naltrindole, and TAN-452 injections in the stress group |
| 2A | Two-way RM ANOVA, *P* < 0.001 for group (F [1, 7] = 146.7)  time (F [7, 49] = 56.60)  group x time interaction (F [7, 49] = 77.19)  One-way RM ANOVA:  sham: *P* = 0.23, F [7, 49] = 1.39  stress: *P* < 0.001, F [7, 49] = 163.5 | Bonferroni for multiple comparison:    ^***^*P* < 0.001, sham versus stress    ^###^*P* < 0.001, compared with the day -3 baseline threshold within stress group  ^&&&^*P* < 0.001, compared with the values before MRS7299 injection in the stress group |
| 2B | Two-way RM ANOVA, *P* < 0.001 for group (F [1, 9] = 66.0)  time (F [5, 45] = 25.18)  group x time interaction (F [5, 45] = 18.08)  One-way RM ANOVA:  sham: *P* = 0.78, F [5, 20] = 0.49  stress: *P* < 0.001, F [5, 25] = 86.95 | Bonferroni for multiple comparison:  ^*^*P* < 0.05, ^***^*P* < 0.001, sham versus stress  ^###^*P* < 0.001, compared with the day -3 baseline threshold in stress group |
| 2C | Two-way RM ANOVA:  *P* < 0.01 for group (F [1, 9] = 13.55)  *P* < 0.001 for time (F [8, 72] = 19.34)  *P* < 0.001 for group x time interaction (F [8, 72] = 10.89)  One-way RM ANOVA:  sham: *P* = 0.57, F [8, 32] = 0.84  stress: *P* < 0.001, F [8, 40] = 56.96 | Bonferroni for multiple comparison:  ^***^*P* < 0.001, sham versus stress  ^&&&^*P* < 0.001, compared with the values before MNB, Bicuc, SNP and BLS challenges in the stress group |
| 2D | One-way RM ANOVA:  *P* < 0.001, F [5, 45] = 91.42 | ^##^*P* < 0.01, ^###^*P* < 0.001, compared with the day -3 baseline threshold |
| 2E | One-way RM ANOVA:  *P* < 0.001, F [8, 32] = 84.70 | ^&&&^*P* < 0.001, compared with the values before each MRS7299 injection |
| 2F | One-way RM ANOVA:  *P* < 0.001, F [8, 32] = 47.06 | ^&&&^*P* < 0.001, compared with the values before each MRS7299 injection |
| 3A | Two-way RM ANOVA: *P* < 0.001 for  group (F [1, 10] = 60.18)  time (F [7, 70] = 11.88)  group x time interaction (F [7, 70] = 7.83)  One-way RM ANOVA:  sham: P = 0.41, F [7, 21] = 0.41  stress: P < 0.001, F [7, 49] = 32.20 | Bonferroni for multiple comparison:  ^***^*P* < 0.001, sham versus stress  ^###^*P* < 0.001, compared with the day -3 baseline threshold in stress group  ^&&&^*P* < 0.001, compared with the values before PGE_2_ injection in the stress group |
| 3B | Two-way RM ANOVA: *P* < 0.001 for  group (F [1, 10] = 22.27)  time (F [6, 60] = 19.81)  group x time interaction (F [6, 60] = 17.65)  One-way RM ANOVA:  sham: *P* = 0.19, F [6,30] = 1.56  stress: *P* < 0.001, F [6,30] = 25.56 | Bonferroni for multiple comparison:  ^***^*P* < 0.001, sham versus stress  ^###^*P* < 0.001, compared with the day -3 baseline threshold in stress group  ^&&&^*P* < 0.001, compared with the values before nalox injection in the stress group |
| 3C | Two-way RM ANOVA,  *P* < 0.01 for group (F [1, 10] = 15.62)  *P <* 0.001 for time (F [6, 60] = 17.49)  *P <* 0.001 for group x time interaction (F [6, 60] = 15.51)  One-way RM ANOVA:  sham: *P* = 0.24, F [6,30] = 1.41  stress: *P <* 0.01, F [6,30] = 31.72 | Bonferroni for multiple comparison:  ^***^*P* < 0.001, sham versus stress  ^###^*P* < 0.001, compared with the day -4 baseline threshold in stress group  ^&&&^*P* < 0.001, compared with the values before nalox injection in the stress group |
| 3D | Two-way RM ANOVA, *P* < 0.001 for  group (F [1, 10] = 45.09)  time (F [6, 60] = 16.68)  group x time interaction (F [6, 60] = 14.15)  One-way RM ANOVA:  sham: *P* = 0.88, F [6,30] = 0.39  stress: *P* < 0.001, F [6,30] = 25.68 | Bonferroni for multiple comparison:    ^***^*P* < 0.001, sham versus stress    ^###^*P* < 0.001, compared with the day -3 baseline threshold in stress group    ^&&&^*P* < 0.001, compared with the values before MNB and Bicuc injections in the stress group |
| 3E | Two-way RM ANOVA, *P* < 0.01 for  group (F [1, 10] = 65.49)  time (F [6, 60] = 23.47)  group x time interaction (F [6, 60] = 20.07)  One-way RM ANOVA:  sham: *P* = 0.87, F [6,30] = 0.40  stress: *P <* 0.001, F [6,30] = 51.87 | Bonferroni for multiple comparison:  ^***^*P* < 0.001, sham versus stress    ^###^*P* < 0.001, compared with the day -4 baseline threshold in stress group  ^&&&^*P* < 0.001, compared with the values before MNB and Bicuc injections in the stress group |
| 4B | Two-way RM ANOVA:  *P* = 0.83 for group (F [2, 13] = 0.19)  *P* < 0.001 for time (F [4, 52] = 185)  *P* = 0.69 for group x time interaction (F [8, 52] = 0.69)  One-way RM ANOVA: *P* < 0.001 for saline (F [4, 12] = 48.88)  BIBN-pre (F [4, 20] = 70.58)  BIBN-post (F [4, 20] = 75.37) | Bonferroni for multiple comparison:  ^###^*P* < 0.001, compared with the day -3 baseline threshold within individual groups  ^!!!^*P* < 0.001, compared with the values before SNP injection within individual groups |
| 6A | Two-way RM ANOVA, *P* < 0.001 for  group (F [1, 10] = 113.6)  time (F [13, 130] = 105.4)  group x time interaction (F [13, 130] = 36.40)  One-way RM ANOVA:  saline: *P <* 0.001, F [13,52] = 45.56  DT: *P <* 0.001, F [13,78] = 108.8 | Bonferroni for multiple comparison:  ^***^*P* < 0.001, stress versus DT  ^###^*P* < 0.001, compared with the day -3 baseline threshold in the saline group  ^&&&^*P* < 0.001, compared with the values before SNP injection in the saline group  ^!!!^*P* < 0.001, compared with the day -3 baseline threshold in the DT group  ^$$$^*P* < 0.001, compared with the values before SNP injection in the DT group |
| 6B | Two-way RM ANOVA,  *P* = 0.04 for group (F [1, 10] = 5.63)  *P <* 0.001 for time (F [20, 20] = 27.32)  *P* = 0.02 for group x time interaction (F [2, 20] = 5.19)  One-way RM ANOVA:  saline: *P <* 0.001, F [2,8] = 57.71  DT: *P <* 0.001, F [2,12] = 15.73 | Bonferroni for multiple comparison:  ^***^*P* < 0.001, as indicated in the figure |
| 6C | Two-way RM ANOVA,  *P* = 0.84 for group (F [10, 100] = 0.04)  *P <* 0.001 for time (F [10, 100] = 142.8)  *P* = 0.86 for group x time interaction (F [10, 100] = 0.54)  One-way RM ANOVA:  saline: *P <* 0.001, F [10,50] = 72.95  DT: *P <* 0.001, F [10,50] = 70.23 | Bonferroni for multiple comparison:  ^###^*P* < 0.001, compared with day -5 thresholds within the saline and DT groups  ^&&&^*P* < 0.001, compared with the values before PGE_2_ injection within the saline and DT groups |
| 6D | Two-way RM ANOVA,  *P* = 0.81 for group (F [1, 10] = 0.05)  *P <* 0.001 for time (F [2, 20] = 40.1)  *P* = 0.66 for group x time interaction (F [2, 20] = 0.41)  One-way RM ANOVA:  saline: *P <* 0.01, F [2,10] = 11.34  DT: *P <* 0.001, F [2,10] = 42.86 | Bonferroni for multiple comparison:  ^&&^*P* < 0.01, compared with the values before PGE_2_ injection within the saline and DT groups |
| 6E | Two-way RM ANOVA,  *P* = 0.77 for group (F [1, 9] = 0.09)  *P <* 0.001 for time (F [10, 90] = 73.45)  *P* = 0.94 for group x time interaction (F [10, 90] = 0.40)  One-way RM ANOVA:  saline: *P <* 0.001, F [10,40] = 55.87  DT: *P <* 0.001, F [10,50] = 31.05 | Bonferroni for multiple comparison:  ^###^*P* < 0.001, compared with day -5 thresholds within the saline and DT groups  ^&&&^*P* < 0.001, compared with the values before PGE_2_ injection within the saline and DT groups |
| 6F | Two-way RM ANOVA,  *P* = 0.96 for group (F [1, 9] = 0.002)  *P <* 0.001 for time (F [2, 18] = 37.4)  *P* = 0.70 for group x time interaction (F [2, 18] = 0.35)  One-way RM ANOVA:  saline: *P <* 0.05, F [2,8] = 8.07  DT: *P <* 0.001, F [2,10] = 61.32 | Bonferroni for multiple comparison:  ^&^*P* < 0.05, compared with the values before PGE_2_ injection within the saline and DT groups |
| 6G | Unpaired two-tailed t-test | *P* < 0.001 |
| 7A | Two-way RM ANOVA, *P* < 0.001 for group (F [1, 9] = 408.2)  time (F [7, 63] = 14.53)  group x time interaction (F [7, 63] = 11.80)  One-way RM ANOVA:  saline: *P* < 0.001, F [7, 28] = 20.23  LD-IL-2: *P* < 0.001, F [7, 35] = 4.68 | Bonferroni for multiple comparison:    ^**^*P* < 0.01, ^***^*P* < 0.001, saline versus LD-IL-2  ^##^*P* < 0.01, ^###^*P* < 0.001, compared with the day -3 baseline threshold in saline group  ^&&&^*P* < 0.001, ^^^^^*P* < 0.001, compared with the day 5 values before PGE_2_ injection in saline and LD-IL-2 group, respectively |
| 7B | Two-way RM ANOVA,  *P* < 0.05 for group (F [1, 10] = 7.88)  *P* < 0.001 for time (F [7, 70] = 52.15)  *P* < 0.001 for group x time interaction (F [7, 70] = 5.91)  One-way RM ANOVA:  saline: *P <* 0.001, F [7, 35] = 34.72  LD-IL-2: *P* < 0.001, F [7, 35] = 24.60 | Bonferroni for multiple comparison:  ^**^*P* < 0.01, ^***^*P* < 0.001, saline versus LD-IL-2  ^##^*P* < 0.01, ^###^*P* < 0.001, compared with the day -3 baseline threshold in saline group  ^!^*P* < 0.05, ^!!!^*P* < 0.001, compared with the day -3 baseline threshold in LD-IL-2 group  ^&&&^*P* < 0.001, compared with the values before PGE_2_ injection in saline group |
| 7C | Two-way RM ANOVA, *P* < 0.001 for  group (F [1, 10] = 67.74)  time (F [10, 100] = 77.79)  group x time interaction (F [10, 100] = 16.20)  One-way RM ANOVA:  saline: *P <* 0.001, F [10, 50] = 69.37  LD-IL-2: *P* < 0.001, F [10, 50] = 33.59 | Bonferroni for multiple comparison:  ^***^*P* < 0.001, saline versus LD-IL-2  ^###^*P* < 0.001, ^!!!^*P* < 0.001, compared with the day -3 baseline thresholds in saline and LD-IL-2 groups, respectively.  ^&&&^*P* < 0.001, compared with the values before each PGE_2_ injection in the saline group |
| 7D | Two-way RM ANOVA:  *P* < 0.01 for group (F [1, 10] = 12.46)  *P* < 0.001 for time (F [3, 30] = 46.95)  *P <* 0.001 for group x time interaction (F [3, 30] = 27.40)  One-way RM ANOVA:  saline: *P* < 0.001, F [3, 15] = 72.87  LD-IL-2: *P* = 0.27, F [3, 15] = 1.42 | Bonferroni for multiple comparison:  ^***^*P* < 0.001, saline versus LD-IL-2  ^&&&^*P* < 0.001, compared with the day -4 baseline scores in the saline group |
| 7E | Two-way RM ANOVA, *P* < 0.001 for  group (F [1, 10] = 35.64)  time (F [7, 70] = 64.25)  group x time interaction (F [7, 70] = 16.52)  One-way RM ANOVA:  saline: *P* < 0.001, F [7, 35] = 47.68  LD-IL-2: *P* < 0.001, F [7, 35] = 34.43 | Bonferroni for multiple comparison:  ^***^*P* < 0.001, saline versus LD-IL-2  ^###^*P* < 0.001, ^!!!^*P* < 0.001, compared with the day -4 baseline values in saline and LD-IL-2 groups, respectively  ^&&&^*P* < 0.001, relative to the values before each PGE_2_ injection in saline group |
| 8A | Two-way RM ANOVA,  *P* < 0.01 for group (F [1, 10] = 15.36)  *P* < 0.001 for time (F [9, 90] = 62.48)  *P* < 0.001 for group x time interaction (F [9, 90] = 8.09)  One-way RM ANOVA:  saline: *P* < 0.001, F [9, 45] = 35.12  LD-IL-2: *P* < 0.001, F [9, 45] = 35.49 | Bonferroni for multiple comparison:  ^**^*P* < 0.01, ^***^*P* < 0.001, saline versus LD-IL-2  ^###^*P* < 0.001, ^!!!^*P* < 0.001, compared with the day -3 baseline thresholds in saline and LD-IL-2 groups, respectively  ^&&&^*P* < 0.001, compared with the values before nalox and Bicuc injections in the saline group |
| 8B | Two-way RM ANOVA,  *P* < 0.05 for group (F [1, 10] = 5.57)  *P* < 0.001 for time (F [5, 50] = 5.72)  *P* < 0.01 for group x time interaction (F [5, 50] = 3.98)  One-way RM ANOVA:  saline: *P* < 0.001, F [7, 35] = 8.04  LD-IL-2: *P* < 0.001, F [7, 35] = 4. 90 | Bonferroni for multiple comparison:  ^*^*P* < 0.05, ^**^*P* < 0.01, saline versus LD-IL-2  ^#^*P* < 0.05, ^##^*P* < 0.01, compared with the day -4 baseline scores in the saline group  ^!!!^*P* < 0.001, compared with the day -4 baseline scores in the LD-IL-2 group.  ^&^*P* < 0.05, ^&&^*P* < 0.01, compared with the values before nalox and bicuc injections in the saline group |
| 8C | Two-way RM ANOVA, *P* < 0.001 for  group (F [1, 10] = 25.85)  time (F [9, 90] = 71.98)  group x time interaction (F [9, 90] = 12.68)  One-way RM ANOVA:  saline: *P* < 0.001, F [9, 45] = 58.81  LD-IL-2: *P* < 0.001, F [9, 45] = 31.38 | Bonferroni for multiple comparison:  ^***^*P* < 0.001, saline versus LD-IL-2  ^###^*P* < 0.001, ^!!!^*P* < 0.001, compared with the day -3 baseline thresholds in saline and LD-IL-2 groups, respectively  ^&&&^*P* < 0.001, compared with the values before nalox and bicuc injections in the saline group |
| 8D | Two-way RM ANOVA,  *P* < 0.01 for group (F [1, 10] = 19.85)  *P* < 0.001 for time (F [7, 70] = 47.10)  *P* < 0.001 for group x time interaction (F [7, 70] = 12.45)  One-way RM ANOVA:  saline: *P* < 0.001, F [7, 35] = 38.54  LD-IL-2: *P* < 0.001, F [7, 35] = 23.66 | Bonferroni for multiple comparison:  ^***^*P* < 0.001, saline versus LD-IL-2  ^###^*P* < 0.001, ^!!!^*P* < 0.001, compared with the day -4 baseline scores in saline and LD-IL-2 groups, respectively  ^&&&^*P* < 0.001, compared with the values before nalox and Bicuc injections in the saline group |
| 9A | Two-way RM ANOVA, *P* < 0.001 for group (F [1, 18] = 192.6)  time (F [5, 90] = 214.9)  group x time interaction (F [5, 90] = 29.36)  One-way RM ANOVA:  saline: *P* < 0.001, F [5, 45] = 107.1  LD-IL-2: *P* < 0.001, F [5, 45] = 140.7 | Bonferroni for multiple comparison:  ^***^*P* < 0.001, saline versus LD-IL-2  ^###^*P* < 0.001, ^!!!^*P* < 0.001, compared with the day -3 baseline thresholds in saline and LD-IL-2 groups, respectively |
| 9B | Two-way RM ANOVA, *P* < 0.001 for group (F [1, 18] = 173.6)  time (F [10, 180] = 50.68)  group x time interaction (F [10, 180] = 48.19)  One-way RM ANOVA:  saline: *P* < 0.001, F [10, 90] = 184.8  LD-IL-2: *P* = 0.11, F [10, 90] = 1.64 | Bonferroni for multiple comparison:  ^***^*P* < 0.001, saline versus LD-IL-2  ^&&&^*P* < 0.001, compared with the values before SNP, MNB, Bicuc, BLS and Sac injections in the saline group |
| 9C | Two-way RM ANOVA,  *P* < 0.01 for group (F [1, 13] = 9.65)  *P* < 0.01 for time (F [1, 13] = 14.32)  *P* < 0.001 for group x time interaction (F [1, 13] = 35.16) | Bonferroni for multiple comparison:  ^***^*P* < 0.001 |
| 9D | Two-way RM ANOVA,  *P* < 0.05 for group (F [1, 9] = 6.29)  *P* = 0.08 for time (F [1, 9] = 3.89)  *P* = 0.05 for group x time interaction (F [1, 9] = 4.98) | Bonferroni for multiple comparison:  ^*^*P* < 0.05, ^**^*P* < 0.01 |
| 9E | Two-way RM ANOVA, *P* < 0.001 for  group (F [1, 14] = 86.71)  time (F [10, 140] = 75.78)  group x time interaction (F [10, 140] = 21.65)  One-way RM ANOVA:  saline: *P* < 0.001, F [10, 70] = 47.43  LD-IL-2: *P* < 0.001, F [10, 70] = 50.05 | Bonferroni for multiple comparison:  ^**^*P* < 0.01, ^***^*P* < 0.001, saline versus LD-IL-2  ^###^*P* < 0.001, ^!!!^*P* < 0.001, compared with the day -3 baseline thresholds in saline and LD-IL-2 groups, respectively  ^&&&^*P* < 0.001, compared with the values before AM281 and AM630 administration in the saline group |
| 9F | Two-way RM ANOVA, *P* < 0.001 for group (F [1, 18] = 25.10)  time (F [8, 144] = 68.39)  group x time interaction (F [8, 144] = 23.10)  One-way RM ANOVA:  saline: *P* < 0.001, F [8, 72] = 143.5  LD-IL-2: *P* < 0.001, F [8, 72] = 16.84 | Bonferroni for multiple comparison:    ^***^*P* < 0.001, saline versus LD-IL-2  ^###^*P* < 0.001, ^!!!^*P* < 0.001, compared with the day -3 baseline thresholds in saline and LD-IL-2 groups, respectively  ^&&&^*P* < 0.001, compared with the values before SNP, MNB and BLS challenges in the saline group |
| 10A | Two-way RM ANOVA,  *P* < 0.05 for group (F [1, 17] = 5.17)  *P* < 0.001 for time (F [5, 85] = 74.69)  *P* < 0.001 for group x time interaction (F [5, 85] = 5.05)  One-way RM ANOVA:  saline: *P* < 0.001, F [5, 45] = 54.90  LD-IL-2: *P* < 0.001, F [5, 40] = 29.04 | Bonferroni for multiple comparison:  ^**^*P* < 0.01, ^***^*P* < 0.001 saline versus LD-IL-2  ^###^*P* < 0.001, ^!!!^*P* < 0.001, compared with the day -3 baseline thresholds in saline and LD-IL-2 groups, respectively |
| 10B | Two-way RM ANOVA, *P* < 0.001 for  group (F [1, 17] = 216.5)  time (F [6, 102] = 28.59)  group x time interaction (F [6, 102] = 39.29)  One-way RM ANOVA:  saline: *P* < 0.001, F [6, 54] = 104.7  LD-IL-2: *P* < 0.001, F [6, 48] = 0.70 | Bonferroni for multiple comparison:  ^***^*P* < 0.001, saline versus LD-IL-2  ^&&&^*P* < 0.001, compared with the values before SNP, MNB and BLS challenges in the saline group |
| 10C | Two-way RM ANOVA,  *P* = 0.37 for group (F [1, 17] = 0.83)  *P* < 0.001 for time (F [2, 34] = 59.10)  *P* = 0.95 for group x time interaction (F [2, 34] = 0.05) | Bonferroni for multiple comparison:  ^***^*P* < 0.001 |
| 10D | Two-way RM ANOVA,  *P* < 0.05 for group (F [1, 17] = 6.51)  *P* < 0.001 for time (F [2, 34] = 32.28)  *P* < 0.001 for group x time interaction (F [2, 34] = 15.90) | Bonferroni for multiple comparison:  ^***^*P* < 0.001 |
| 11A | Two-way RM ANOVA, *P* < 0.001 for  group (F [1, 13] = 97.95)  time (F [9, 117] = 112.8)  group x time interaction (F [9, 117] = 34.29)  One-way RM ANOVA:  saline: *P* < 0.001, F [9, 63] = 55.25  DT: *P* < 0.001, F [9.54] = 133.4 | Bonferroni for multiple comparison:  ^**^*P* < 0.01, ^***^*P* < 0.001, saline versus DT  ^###^*P* < 0.001, ^!!!^*P* < 0.001, compared with the day -3 baseline thresholds within the saline and DT groups, respectively  ^&&&^*P* < 0.001, compared with the day 16 thresholds before SNP injection in the DT group |
| 11B | Two-way RM ANOVA, *P* < 0.001 for  group (F [1, 14] = 28.66)  time (F [8, 112] = 185.5)  group x time interaction (F [8, 112] = 18.57)  One-way RM ANOVA:  saline: *P* < 0.001, F [8, 56] = 107.7  DT: *P* < 0.001, F [8, 56] = 97.03 | Bonferroni for multiple comparison:  ^***^*P* < 0.001, saline versus DT  ^###^*P* < 0.001, ^!!!^*P* < 0.001, compared with the day -3 baseline thresholds within the saline and DT groups, respectively  ^&&&^*P* < 0.001, compared with the day 8 thresholds before PGE_2_ injection in the DT group |
| 11C | Two-way RM ANOVA, *P* < 0.001 for  group (F [1, 14] = 101.3)  time (F [7, 98] = 153.6)  group x time interaction (F [7, 98] = 25.87)  One-way RM ANOVA:  saline: *P* < 0.001, F [7, 49] = 151.1  LD-IL-2: *P* < 0.001, F [7, 49] = 58.48 | Bonferroni for multiple comparison:  ^***^*P* < 0.001, saline versus LD-IL-2  ^###^*P* < 0.001, ^!!!^*P* < 0.001, compared with the day -3 baseline thresholds in saline and LD-IL-2 groups, respectively  ^&&&^*P* < 0.001, compared with the values before naloxone in the saline group |
| 11D | All groups:  Two-way RM ANOVA, *P* < 0.001 for  group (F [3, 16] = 26.43)  time (F [6, 96] = 178.5)  group x time interaction (F [18, 96] = 12.6)  LD-IL-2 vs LD-IL-2+naloxone:  Two-way RM ANOVA,  *P* = 0.12 for group (F [1, 8] = 2.95)  *P* < 0.001 for time (F [6, 48] = 79.76)  *P* = 0.12 for group x time interaction (F [6, 48] = 1.07)  One-way RM ANOVA:  saline: *P* < 0.001, F [6, 24] = 75.09  LD-IL-2: *P* < 0.001, F [6, 24] = 40.24  LD-IL-2+naloxone: *P* < 0.001, F [6, 24] = 40.58  Naloxone: *P* < 0.001, F [6, 24] = 104.1 | Bonferroni for multiple comparison:  Day 5, *P* < 0.01, day 7, *P* < 0.001, saline versus LD-IL-2  Day 7, *P* < 0.001, saline versus LD-IL-2+naloxone  Day 9, *P* < 0.001, saline versus naloxone  Day 7 and 9, *P* < 0.001, LD-IL-2+naloxone versus naloxone  ^###^*P* < 0.001, ^!!!^*P* < 0.001, ^+++^*P* < 0.001, ^^^^^*P* < 0.001, compared with the day -3 baseline thresholds in saline, LD-IL-2, naloxone and LD-IL-2+naloxone groups, respectively. |
| 11E | Two-way ANOVA,  *P* < 0.001 for LD-IL-2 effects, (F [1, 16] = 85.66)  *P* < 0.01 for naloxone effects, (F [1, 16] = 11.91)  *P* = 0.58 for interaction, (F [1, 16] = 0.32 | Bonferroni for multiple comparison:  ^***^*P* < 0.001, saline versus LD-IL-2 and naloxone versus LD-IL-2+naloxone  *P* = 0.07, saline versus naloxone  *P* = 0.35, LD-IL-2 versus LD-IL-2+naloxone |
| 12A | Two-way RM ANOVA, *P* < 0.001 for  group (F [2, 15] = 84.79)  time (F [10, 150] = 39.60)  group x time interaction (F [20, 150] = 8.08)  One-way RM ANOVA:  stress+IgG: *P* < 0.001, F [11, 55] = 28.14  stress+anti-IFNγ: *P* < 0.001, F [11, 55] = 23.81 | Bonferroni for multiple comparison:  Day 1-6, 13”,15” and 17”, *P* < 0.001, sham versus stress+IgG and stress+anti-IFNγ  Day 1, *P* < 0.05, stress+IgG versus stress+anti-IFNγ  ^###^*P* < 0.001, compared with the day -5 baseline thresholds within the stress+IgG and stress+anti-IFNγ groups  ^&&&^*P* < 0.001, compared with the values before SNP, BLS or nalox challenges within the stress+IgG and stress+anti-IFNγ groups |
| 12B | Two-way RM ANOVA,  *P* = 0.05 for group (F [1, 10] = 5.03)  *P* < 0.001 for time (F [12, 120] = 60.74)  *P* = 0.62 for group x time interaction (F [12, 120] = 0.82)  One-way RM ANOVA:  stress+IgG: *P* < 0.001, F [12, 36] = 26.67  stress+anti-IL-10: *P* < 0.001, F [12, 84] = 41.34 | Bonferroni for multiple comparison:  ^###^*P* < 0.001, compared with the day -5 baseline thresholds within the stress+IgG and stress+anti-IL-10 groups  ^&&&^*P* < 0.001, compared with the values before SNP, BLS or nalox challenges within the stress+IgG and stress+anti-IL-10 groups |
| 12C | One-way RM ANOVA:  stress+anti-TGFβ: *P* < 0.001, F [16, 96] = 23.68 | Bonferroni for multiple comparison:  ^###^*P* < 0.001, ^&&&^*P* < 0.001, compared with the day -5 baseline thresholds and the day 27 thresholds before SNP injection, respectively |
| 13A | Two-way RM ANOVA,  *P* < 0.01 for group (F [2, 11] = 7.94)  *P* < 0.001 for time (F [10, 110] = 44.62)  *P* < 0.001 for group x time interaction (F [20, 110] = 6.02)  One-way RM ANOVA:  stress+IgG: *P* < 0.001, F [10, 30] = 6.48  stress+anti-IFNγ: *P* < 0.001, F [10, 40] = 31.11  stress+anti-IL-10: *P* < 0.001, F [10, 40] = 42.16 | Bonferroni for multiple comparison:  ^***^*P* < 0.001, stress+IgG versus stress+anti-IL-10 and stress+anti-IFNγ  ^###^*P* < 0.001, compared with the day -3 baseline thresholds within the stress+anti-IL-10 and stress+anti-IFNγ groups  ^!!^*P* < 0.01, compared with the day -3 baseline thresholds in stress+IgG group  ^&&&^*P* < 0.001, compared with the values before SNP, BLS and nalox challenges within the stress+anti-IFNγ and stress+anti-IL-10 groups |
| 13B | One-way RM ANOVA:  stress+anti-TGFβ: *P* < 0.001, F [11, 44] = 26.82 | Bonferroni for multiple comparison:  ^###^*P* < 0.001, compared with the day -3 baseline thresholds  ^&&&^*P* < 0.001, compared with the day 9 thresholds before SNP injection |
| 13C | One-way RM ANOVA:  *P* < 0.001, F [16, 96] = 140.8 | Bonferroni for multiple comparison:  ^###^*P* < 0.001, compared with the day -3 baseline thresholds  ^&&&^*P* < 0.001, compared with the day 27 thresholds before the SNP injection |
| 13D | One-way RM ANOVA:  *P* < 0.001, F [16, 96] = 139.2 | Bonferroni for multiple comparison:  ^!!!^*P* < 0.001, compared with the day -3 baseline thresholds  ^&&&^*P* < 0.001, compared with the day 9 thresholds before the SNP injection |
| 14A | Two-way RM ANOVA, *P* < 0.001 for  group (F [2, 14] = 38.23)  time (F [11, 154] = 96.47)  group x time interaction (F [22, 154] = 15.19)  One-way RM ANOVA:  control IgG: *P* < 0.001, F [11, 44] = 56.59  anti-CGRP: *P* < 0.001, F [11, 66] = 57.93  LD-IL-2: *P* < 0.001, F [11, 44] = 24.63 | Bonferroni for multiple comparison:  ^***^*P* < 0.001, LD-IL-2 versus anti-CGRP and control IgG  ^^^^^*P* < 0.001, LD-IL-2 versus control IgG  ^###^*P* < 0.001, compared with the day -3 baseline thresholds in control IgG and anti-CGRP groups  ^!!!^*P* < 0.001, compared with the day -3 baseline thresholds in LD-IL-2 group  ^&&&^*P* < 0.001, ^+++^*P* < 0.001, compared with the values before SNP, Bicuc, or BLS challenges within the control IgG and anti-CGRP groups, respectively |
| S1 | One-way RM ANOVA:  saline: *P* < 0.001, F [10, 50] = 69.37 | Bonferroni for multiple comparison:  ^###^*P* < 0.001, compared with the pre-stress baseline threshold  ^&&&^*P* < 0.001, compared with the values before each PGE_2_ injection |
